# Single-cell transcriptomic and genomic changes in the aging human brain

**DOI:** 10.1101/2023.11.07.566050

**Authors:** Ailsa M. Jeffries, Tianxiong Yu, Jennifer S. Ziegenfuss, Allie K. Tolles, Yerin Kim, Zhiping Weng, Michael A. Lodato

## Abstract

Aging brings dysregulation of various processes across organs and tissues, often stemming from stochastic damage to individual cells over time. Here, we used a combination of single-nucleus RNA-sequencing and single-cell whole-genome sequencing to identify transcriptomic and genomic changes in the prefrontal cortex of the human brain across life span, from infancy to centenarian. We identified infant-specific cell clusters enriched for the expression of neurodevelopmental genes, and a common down-regulation of cell-essential homeostatic genes that function in ribosomes, transport, and metabolism during aging across cell types. Conversely, expression of neuron-specific genes generally remains stable throughout life. We observed a decrease in specific DNA repair genes in aging, including genes implicated in generating brain somatic mutations as indicated by mutation signature analysis. Furthermore, we detected gene-length-specific somatic mutation rates that shape the transcriptomic landscape of the aged human brain. These findings elucidate critical aspects of human brain aging, shedding light on transcriptomic and genomics dynamics.

## Main Text

Aging is a universal, multifaceted biological process that affects all the systems in the body^1^. Bulk RNA-sequencing studies of aging have revealed disruptions to essential cellular processes such as transcription, translation, and growth factor signaling^2,3^, with processes involved in mitochondrial function, neuronal activity, and DNA damage being dysregulated in the aging brain^4,5^. Cell-type-specific changes during aging are obscured in bulk analysis and are poorly understood. This is a major knowledge gap in the human brain, where molecularly distinct cell types carry out specific functions throughout life. Single-cell whole-genome sequencing (scWGS) and other high-resolution techniques have demonstrated that somatic mutations accumulate in human neurons during aging and in age-related disease, raising the possibility that such variants could contribute to transcriptional dysregulation and the concomitant increased susceptibility dysfunction and disease that accompanies advanced age^6–11^. The advent of single-cell genomic technologies has provided high resolution in both DNA and RNA sequencing. Single-cell and single-nucleus RNA-sequencing (snRNA-seq) analysis has identified age-related changes in several organs^12^, and snRNA-seq of the human brain has identified several transcriptional states^13–15^ that are disrupted in age-associated disease^16,17^. However, the single-cell transcriptional changes associated with non-pathological aging that set the stage for diseases of brain aging remain incompletely defined.

To better understand the dynamics of human brain aging, we generated droplet-based snRNA-seq^18^ libraries of fresh-frozen human pre-frontal cortex (PFC) (Fig. 1a) from 13 donors of different ages, ranging from infant to centenarian (Table 1 and Table S1). 290,814 nuclei remained after quality control and artifact filtering^19^, a mean of 22,370/donor (fig. S1). Dimensionality reduction and hierarchical clustering^20^ of these nuclei yielded 32 clusters (Fig. 1b). We annotated these clusters with a previously published human PFC data set^21^ as a reference (Fig 1c, Table S2) and as expected we recovered excitatory neurons from various cortical layers, four subtypes of inhibitory neurons (IN-PV, IN-SST, IN-SV2C, IN-VIP), microglia, oligodendrocytes, oligodendrocyte precursor cells (OPCs), astrocytes and endothelial cells. Expression of canonical marker genes for each cell type is cluster-specific (fig. S2). Within these broad classes, we identified sub-classes of cells that despite being similar to their cells in the same class, populated distinct clusters (Fig. 1d). Excitatory neurons expressed more than twice as many genes on average as glial and endothelial cells (Fig. 1d).

**Fig 1.**
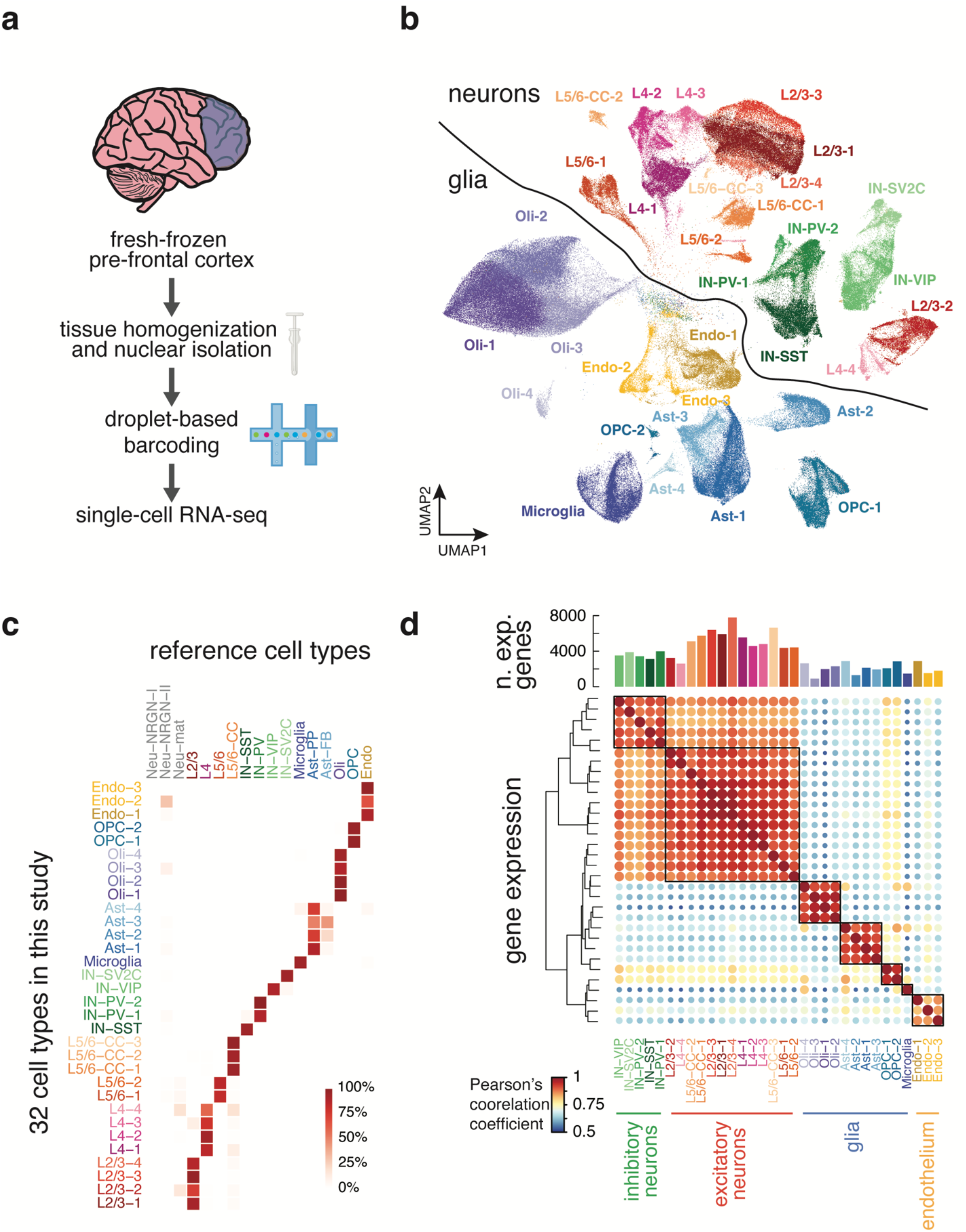
Droplet-based snRNA-seq of human PFC and cell-type classification. (**a**) Experimental schematic of droplet-based snRNA-seq. (**b**) Dimensional reduction and clustering of all cells post-filtration yielded multiple clusters for each cell type (Oli, oligodendrocyte; AST, astrocyte; Endo, endothelial). (**c**) Percentage of cells in each cluster of our data that correspond to the annotated reference cluster. (**d**) Correlation of gene expression profiles for each sub-cluster within a cell type correspond most closely to the cells of the same lineage based on Pearson’s correlation coefficient. Bar plot above heatmap shows the number of genes expressed in each cluster.

**Table 1.**
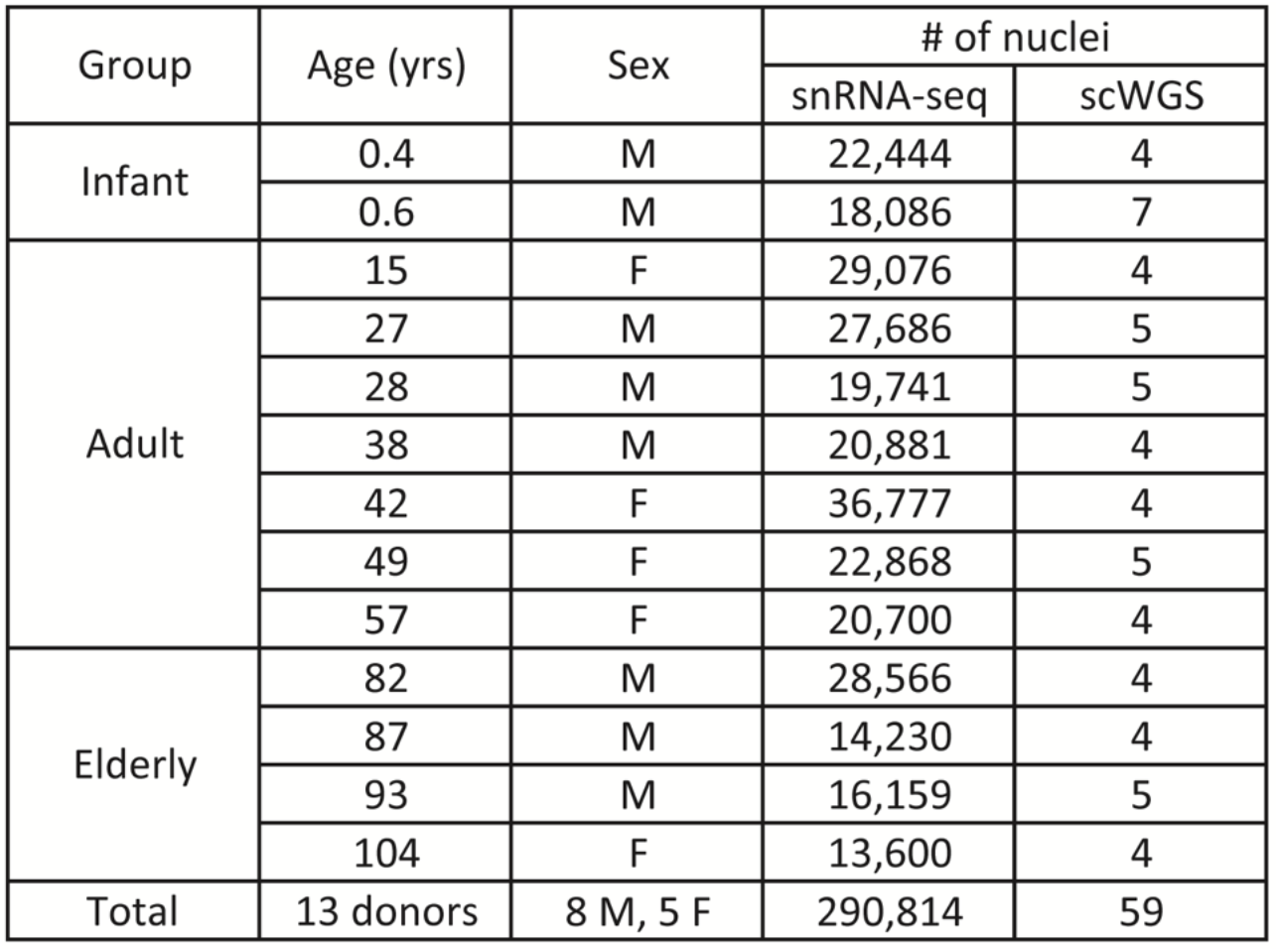
Sample information for single-nucleus RNA-sequencing and single-cell whole genome sequencing. Group, age, and sex information for each donor brain. Number of unsorted nuclei input for snRNA-seq as well sorted NeuN+ neuronal nuclei used for scWGS from each brain. Dorsolateral prefrontal cortex was used for all experiments.

Age-associated neurological diseases such as Alzheimer’s disease (AD), Parkinson’s disease (PD), and cancer result in the loss or expansion of specific brain cell types in the elderly brain. We found no difference in the overall ratios of neurons to glia or excitatory neurons to inhibitory neurons during non-pathological aging, nor did we observe evidence of the expansion of reactive microglia in the elderly brain (Extended Data Fig. 1, Table S3). However, we did identify sub-clusters in L2/3 neurons, L4 neurons, and astrocytes exclusively or nearly exclusively comprising infant cells (Fig. 2a). Infant donors contributed no L2/3 or L4 neurons to other neuronal cell clusters, but some infant astrocytes clustered with adult and elderly astrocytes. Infant-specific clusters were distinguished by the expression of neurodevelopmental genes (Fig. 2b and Table S4 and S5), suggesting the transcriptomes of some infant neurons and astrocytes are still maturing in infancy. Our re-analysis of a published snRNA-seq dataset of human PFC comprising fetal development through adulthood^22^ confirmed the patterns of down-and upregulated genes we observed in our dataset (Extended Data Fig. 2 and fig. S3). Finally, the abundance of OPCs decreased during aging (p-value = 2.56 x 10^-2^, Wilcoxon Rank-sum test), being highest in infant donors and decreasing over lifespan (Fig. 2c), while mature oligodendrocytes increased during aging in the brain (p-value = 2.56 x 10^-2^, Wilcoxon Rank-sum test comparing infant to adult and elderly). These data suggest that the pool of OPCs differentiates into mature oligodendrocytes during life with incomplete replacement, thus the capacity to generate new oligodendrocytes may diminish in elderly humans.

**Fig. 2.**
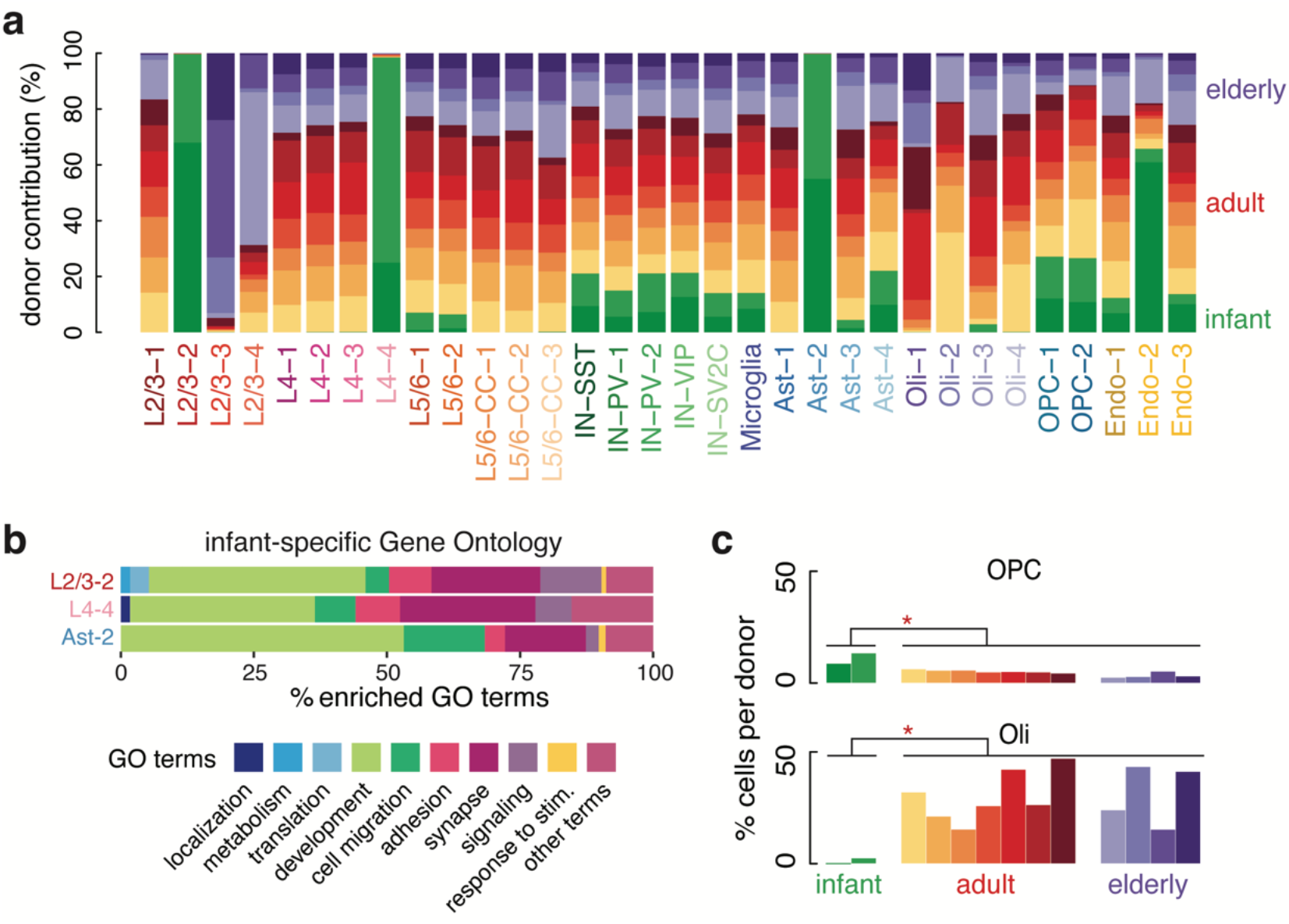
Transcriptionally unique excitatory neurons and astrocytes detected in infant PFC. (**a**) Clusters plotted by donor contribution as a percentage of total cells in the cluster. L2/3-2, L4-4, and AST-2 are infant-specific. (**b**) GO terms derived from differentially expressed genes up-regulated in infant-specific clusters plotted as general categories (see Table S5 for full term list and category designation). Development related terms are most common. (**c**) Contribution of OPCs (top) and oligodendrocytes (bottom) to the total cells in each donor, plotted as a percentage. OPCs are most abundant in infants and their population declines later in life. Oligodendrocytes are most abundant in adulthood and nearly absent in infancy.

Differential expression analysis by cell type comparing the four elderly cases with the seven adult cases yielded 607 genes that change significantly with age (Log_2_(elderly/adult) magnitude > 0.5, p-value < 0.05) (Fig. 3b). In every cell type, there were more genes down-regulated in aging than up-regulated (Wilcoxon Signed-rank test, p-value = 1.66 x 10^-3^) (Fig. 3a). On average non-neuronal cells had a larger number of genes up-regulated in response to aging than neuronal cells (Wilcoxon Rank-sum test, p-value = 4.26 x 10^-3^). Microglia had the most up- and down-regulated genes (117 and 159, respectively) of all cell types. 30 genes down-regulated in aging were commonly downregulated across multiple types of brain cells (Table S6) (p<0.001, random permutation test). For example, the heat shock protein HSPA8 and the cytoskeletal protein TUBA1A were significantly downregulated in all 13 brain cell types during aging. Other commonly downregulated genes across cell types included additional cytoskeletal genes such as TUBB2A (down in 11/13 cell types), TUBA1B (down in 9/13 cell types), and TUBB3 (down in 6/13 cell types), the calmodulin genes CALM1, CALM2, and CALM3 (down in 5, 8, and 9 of 13 cell types), and the vesicle protein VAMP2 (down in 11/13 cell types). In contrast, only one transcript, a poorly characterized mRNA called AC119673.2, was commonly up-regulated in multiple types of neurons and glia.

**Fig. 3.**
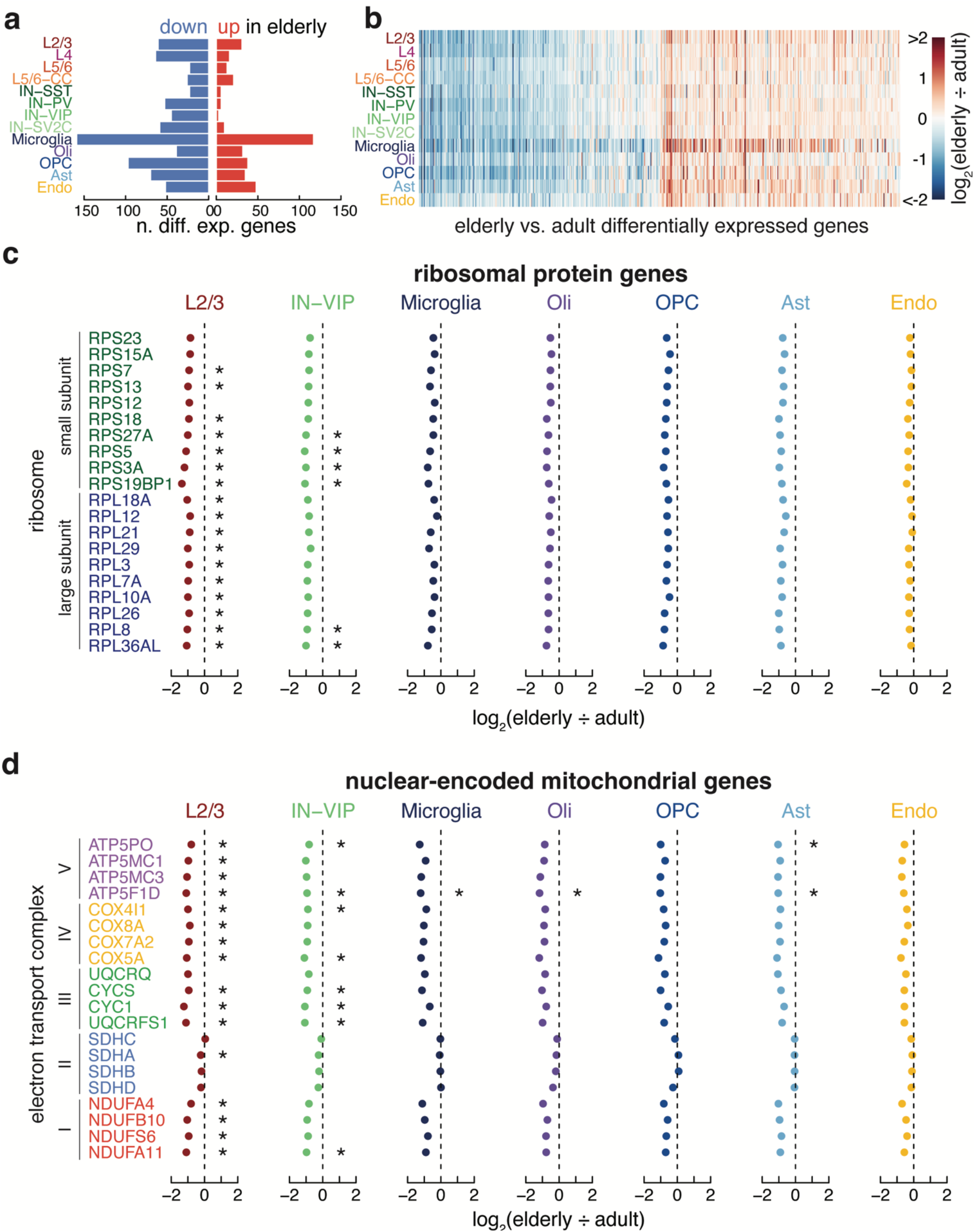
Common down-regulation of genes across cell types. (**a**) The number of down (blue) and up (red) -regulated genes for each cell type. (**b**) Heatmap showing Log_2_ fold-change of elderly vs. adult differentially expressed genes across cell types.(**c**) Log_2_ fold change of elderly vs. adult ribosomal protein genes from both the small and large subunit. (**d**) Log_2_ fold change of elderly vs. adult nuclear encoded mitochondrial electron transport chain genes from all five complexes. (*, p < 0.05, two-sided T-test).

The commonality seen in down-regulated genes across cell types suggested that common pathways were dysregulated across cell types in the aging brain. The ribosomal proteins RPS27A, RPL3, and RPL15 were significantly downregulated during aging in at least one subtype each of excitatory neuron, inhibitory neuron, and glial cell (Table S7), prompting us to inspect the expression level of all ribosomal genes across all brain subtypes. Indeed, we observed a near-universal trend of decreasing expression of every protein from both the small and large ribosomal subunits during aging, far more than expected by chance (p-values - L2/3: 3.92 x 10^-12^; IN-VIP: 4.80 x 10^-14^; oligodendrocytes: 4.29 x 10^-13^; OPCs: 2.49 x 10^-9^; astrocytes: 2.49 x 10^-9^; microglia: 3.27 x 10^-7^; endothelial: 1.04 x 10^-3^, Fisher’s exact test) (Fig. 3c and Extended Data Fig. 3). Along these lines, the nuclear-encoded mitochondrial electron transport chain gene NDUFA4 significantly decreased in excitatory neurons, inhibitory neurons, and multiple types of glia, while ATP5F1D and COX4I1, also key nuclear-encoded mitochondrial electron transport chain genes, decreased in multiple types of aged glia (Table S7). Focused analysis demonstrated universal down-regulation across cell types for most genes in these complexes (p-values – L2/3: 3.49 x 10^-11^; IN-VIP: 4.77 x 10^-13^; oligodendrocytes: 2.13 x 10^-10^; OPCs: 7.71×10^-9^; astrocytes: 1.10 x 10^-9^; microglia: 3.49 x 10^-11^; endothelial: 4.94 x 10^-9^, Fisher’s exact test). Interestingly, Complex II was the only complex to have no significantly down-regulated genes (Fig. 3d and Extended Data Fig. 4).

In accordance with the common downregulation observed in ribosomal proteins and mitochondrial proteins, gene ontology (GO) analysis of downregulated genes yielded common terms across cell types (Fig. 4a and Table S8). Across all groups, terms related to “housekeeping” functions such as translation, metabolism, homeostasis, intracellular localization, and intracellular transport were significantly enriched in the downregulated genes, but most prominently in the neurons. To probe this relationship further, we defined a set of housekeeping genes in our dataset as those genes detected in all brain subtypes, including endothelial cells and microglia which derive from a distinct embryological origin from neurons and other glia, and measured their expression changes during aging. By the same logic, we defined “neuron-specific” genes as those detected in all neuron subtypes but absent in endothelial and microglial cells. Expression of these housekeeping genes decreased in elderly endothelial cells, oligodendrocytes, and all neuron subtypes but L5/6 neurons (Fig. 4b). In contrast, neuron-specific genes did not generally decrease across neuron subtypes during aging, with SST+ inhibitory neurons being the only exception (Extended Data Fig. 5a). Thus, neurons lose expression of genes related to general cell function but maintain cell identity in the aging brain.

**Fig. 4.**
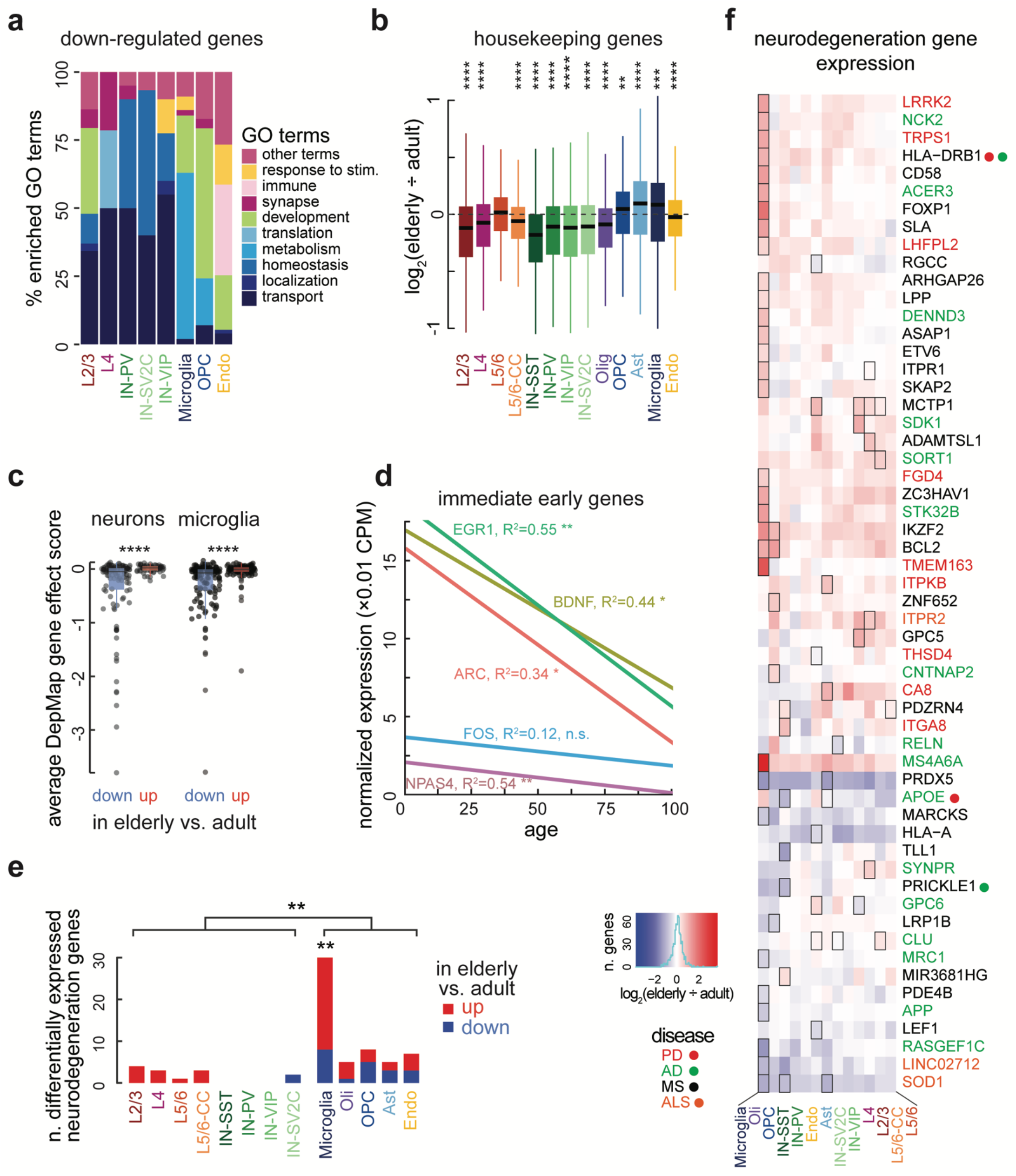
(below) Aging-related down-regulation of essential housekeeping functions in neurons. (**a**) GO terms of genes downregulated in aging plotted as general categories (see Table S8 for full GO results). Housekeeping functions (shades of blue) are most commonly downregulated in neurons and microglia. Only cell types with significantly enriched GO terms shown. (**b**) Housekeeping genes, which are significantly downregulated in elderly brains relative to adults in 7/8 neuron types. Boxes show median, first and third quartiles. Whiskers show one and a half times the interquartile range beyond the first and third quartiles. (**c**) Boxplots showing the mean gene effect score for all the down and up -regulated genes in the DepMap database from the Broad Institute. The down-regulated genes for both neurons (left plot) and microglia (right plot) are more essential than the up-regulated genes (two-sided T-test). Boxes and whiskers formatted as in b. All data points shown. Points beyond whiskers are outliers. (**d**) Expression of immediate early genes ARC, BDNF, EGR1, and NPAS4, in excitatory neurons decreases with age. **(e)** Number of DEGs linked to age-associated neurodegenerative disease by subtype. Red proportions depict up DEGs, blue denotes down DEGs. Microglial DEGs were enriched for neurodegeneration GWAS hits (p = 0.0068, χ^2^ test), and proportionally more glial DEGs were neurodegeneration GWAS hits than neuronal DEGs (7.2% vs. 2.5%, p = 0.0099, two sided t-test). **(f)** Heatmap showing log_2_(FC) of neurodegeneration GWAS hits that were DEGs in at least one subtype. Gene names are colored by disease. Dots near gene names reflect genes with >1 association. (*, p < 0.05; **, p < 0.01; ***, p < 0.001; ****, p < 0.0001) Boxes around cells in heatmap denote significant DEGs.

Public databases of gene essentiality suggest genes downregulated in aging reduce brain cell viability. The DepMap database scores the gene essentiality based on survival rates after knock-out in hundreds of cancer cell lines. Using the DepMap database, we found that downregulated genes were more often essential for cell survival than upregulated genes in neurons and microglia (neurons: p-value = 7.33 x 10^-7^; microglia: p-value = 9.09 x 10^-7^, two-sided T-test) (Fig. 4c). These data indicate that metabolic genes that are necessary for basic cellular functions are downregulated during aging, suggesting cells of the elderly brain are less active. Indeed, the expression of immediate early genes, which are activated rapidly during neuronal stimulation, decreases during brain aging (Fig. 4d and Extended Data Fig. 5b).

Aging is accompanied by an increased incidence of several neurodegenerative diseases, including AD, PD, Amyotrophic Lateral Sclerosis (ALS), Multiple Sclerosis (MS), and others. The way aging influences the progression of these related but distinct diseases in the various cells of the brain is unclear. Our snRNA-seq dataset, which was not presorted to enrich any specific population, allowed us to examine expression changes linked to these diseases in a cell-type-specific manner. Using the NHGRI-EBI Catalog of human genome-wide association studies (GWAS), we defined a set of genes implicated in at least one age-associated neurodegenerative disease, and asked which of these genes are differentially regulated across aging brain cell types (Figure 4e). Surprisingly, more neurodegeneration GWAS hits were differentially expressed in glia than in neurons (p = 7×10^-3^, two-sided T-test), with microglia in particular being enriched for differentially expressed neurodegeneration genes (p = 6.8×10^-3^, χ^2^ test). In contrast to the overall trend of more gene downregulation during aging, neurodegeneration differentially expressed genes tended towards upregulation in aging (p = 7.6×10^-4^, χ^2^ test). Genes that were significantly up- or downregulated in one cell type trended in the same direction across cell types (Figure 4f), suggesting common mechanisms of age-related dysregulation.

Somatic mutations accumulate in cells of the human body during life across many cell types^10,23–27^ including in post-mitotic neurons of the human brain^7,9^. Neuronal somatic mutation rates are higher in transcribed regions of the neuronal genome than transcriptionally silent loci ^7,8,10^, and amongst transcribed genes, bulk brain RNA-seq analysis revealed that those with the highest RNA expression levels showed the highest somatic mutation rate^6,8^. Mutation signature analysis, a set of methods used to infer the molecular mechanism causing mutations by analyzing nucleotide context in which mutations occur, has implicated the activity of several DNA repair pathways in generating somatic mutations in neurons^6,8,11^, suggesting that the expression level of these repair proteins are relevant for controlling the somatic mutation burden. Since the genome is foundational to all cellular programs, the predicted effects of somatic mutation include missense and nonsense mutations in coding regions and dysregulation of gene regulatory networks^28,29^. Thus, the upstream causes and downstream effects of single-cell somatic mutations are both linked to single-cell gene expression.

To compare changes in the neuronal transcriptome to changes to the somatic mutation burden of individual neurons, we performed single-cell whole genome sequencing (scWGS) using primary template-derived amplification (PTA)^8,30^ on neurons from the same brain region and donors analyzed by snRNA-seq (Table S9). We used the SCAN2^7^ algorithm to identify somatic single nucleotide variants (sSNVs) in scWGS data from each sample (Table S10). In agreement with previous reports, our analysis suggested somatic SNVs accumulate at a rate of 15.5 per neuron per year (R^2^ = 0.92, p = 3.21 x 10^-34^) (Extended Data Fig. 6a), with the overall pattern of mutations resembling a known pattern of mutation called SBS5 (cosine similarity = 0.96), first identified by the Catalogue Of Somatic Mutations In Cancer (COSMIC) Consortium, which accumulates during life across many tissues^24,31,32^.

Comparing changes in neuronal gene expression during life to age-related patterns of somatic mutation in neurons revealed relationships between the genome and transcriptome in aging. The overall spectrum of neuron somatic SNVs (Extended Data Fig. 6b) was composed of two distinct signatures, which we name A1 and A2 (Fig. 5a). Signature A1 mutation burden strongly correlates with age of donor (R^2^=0.94, p=2.49 x 10^-37^) (Figure 5b), accounting for 13.8 of the 15.5 mutations per year. Signature A1 burden also correlated strongly with neuronal gene expression levels (Fig 5c, S4), demonstrating the transcription in neurons sensitizes some loci to specific types of somatic mutation, as well as gene expression levels in other cell types (Fig. S4). In line with this, the degree of transcriptional strand bias in somatic SNVs, which results from asymmetric repair of DNA lesions on the template strand at transcribed loci ^33^, increased with excitatory neuron expression levels (Figure 5d), supporting the notion that transcription-coupled DNA repair plays a role in generating neuronal somatic mutations.

**Fig. 5.**
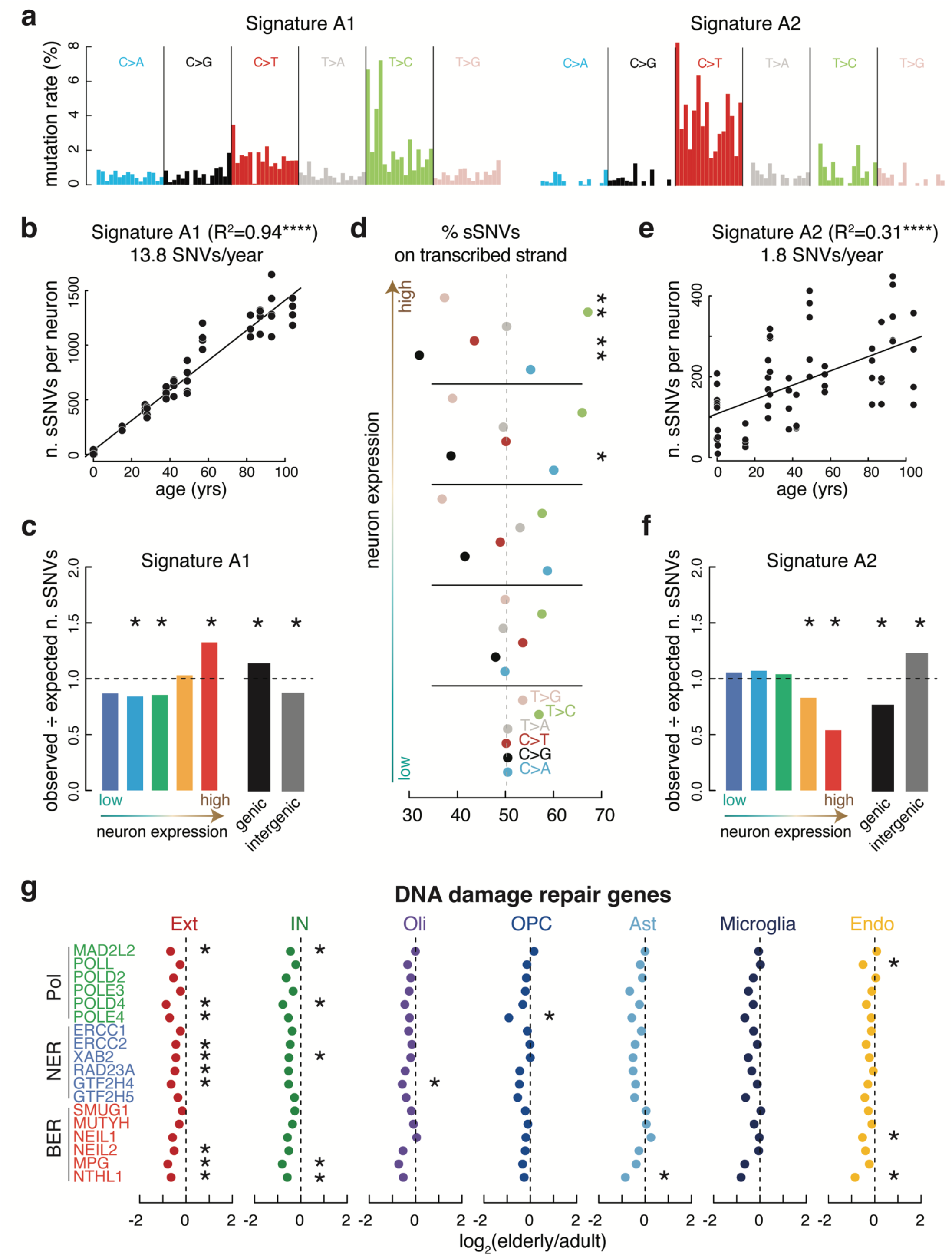
(below) scWGS reveals sSNV mutation signatures linked to expression. (**a**) De novo mutation signature analysis of somatic SNVs (sSNVs) in human neurons revealed two signatures, A1 dominated by T>C mutations and A2 dominated by C>T mutations. Trinucleotide contexts are the same as shown in fig.S13. (**b**) Number of Signature A1 sSNVs in each neuron plotted by age. Signature A1 strongly correlates with age (R^2^=0.94) with an extrapolated mutation rate of 13.8 SNVs/year. (**c**) sSNV enrichment of Signature A1 in coding regions plotted by expression quantile (left) and genic vs. intergenic regions (right). Signature A1 is enriched in the highest expressed genes and genic regions. (**d**) Percent of total sSNVs derived from the transcribed strand broken down by expression quantile. T>C and C>T strand bias increases with expression. (**e**) Number of Signature A2 sSNVs in each neuron plotted by age. Signature A2 correlates with age (R^2^=0.31) with an extrapolated mutation rate of 1.8 SNVs/year. (**f**) sSNV enrichment of Signature A2 in coding regions plotted by expression quantile (left) and genic vs. intergenic regions (right). Signature A2 is depleted in the highest expressed genes and enriched in the lowest expressed genes as well as intergenic regions. (**g**) Log_2_ fold change of elderly/adult expression data from snRNA-seq showing reduced expression in key neuron DNA damage repair pathway proteins (two-sided T-test). Polymerases, green; nucleotide excision repair, blue; base excision repair, red. (*, p < 0.05; ****, p < 0.0001).

Signature A2 accounted for fewer (1.8, R^2^ = 0.31, p = 4.98 x 10^-6^) age-related mutations per year (Figure 5e) and clustered with the known COSMIC signature SBS30 (Extended Data Fig. 6c). SBS30 mutations in tumors are depleted in coding regions (COSMIC Database) and are associated with loss of function mutations in the base excision repair gene NTHL1^34^. Furthermore, NTHL1 knockout in human colon organoids results in increased mutations resembling SBS30^35^. In line with this, Signature A2 mutation rates anticorrelate with neuron gene expression levels and are enriched in intergenic regions (Figure 5f) and NTHL1 is one of the DNA repair genes that decreases in expression in aging neurons (Figure 5g, Extended Data Fig. 7; two-sided T-test), providing a possible mechanism allowing Signature A2 mutations to accumulate during life.

Several recent studies have argued that long genes are downregulated in aging across many organs, presumably because longer genes naturally have a higher chance of acquiring transcription-blocking DNA damage ^36–38^. Surprisingly, downregulated genes were shorter than the average expressed gene across all brain cell types (Figure 6a), with this effect being strongest in excitatory and inhibitory neurons (p = 5.52 x 10^-18^ and 8.42 x 10^-19^ respectively, Wilcoxon rank sum test). Upregulated genes were significantly longer than average in all cell types, with a weaker effect in non-neuronal cell types (Figure 6b, Extended Data Fig. 8).

**Fig. 6.**
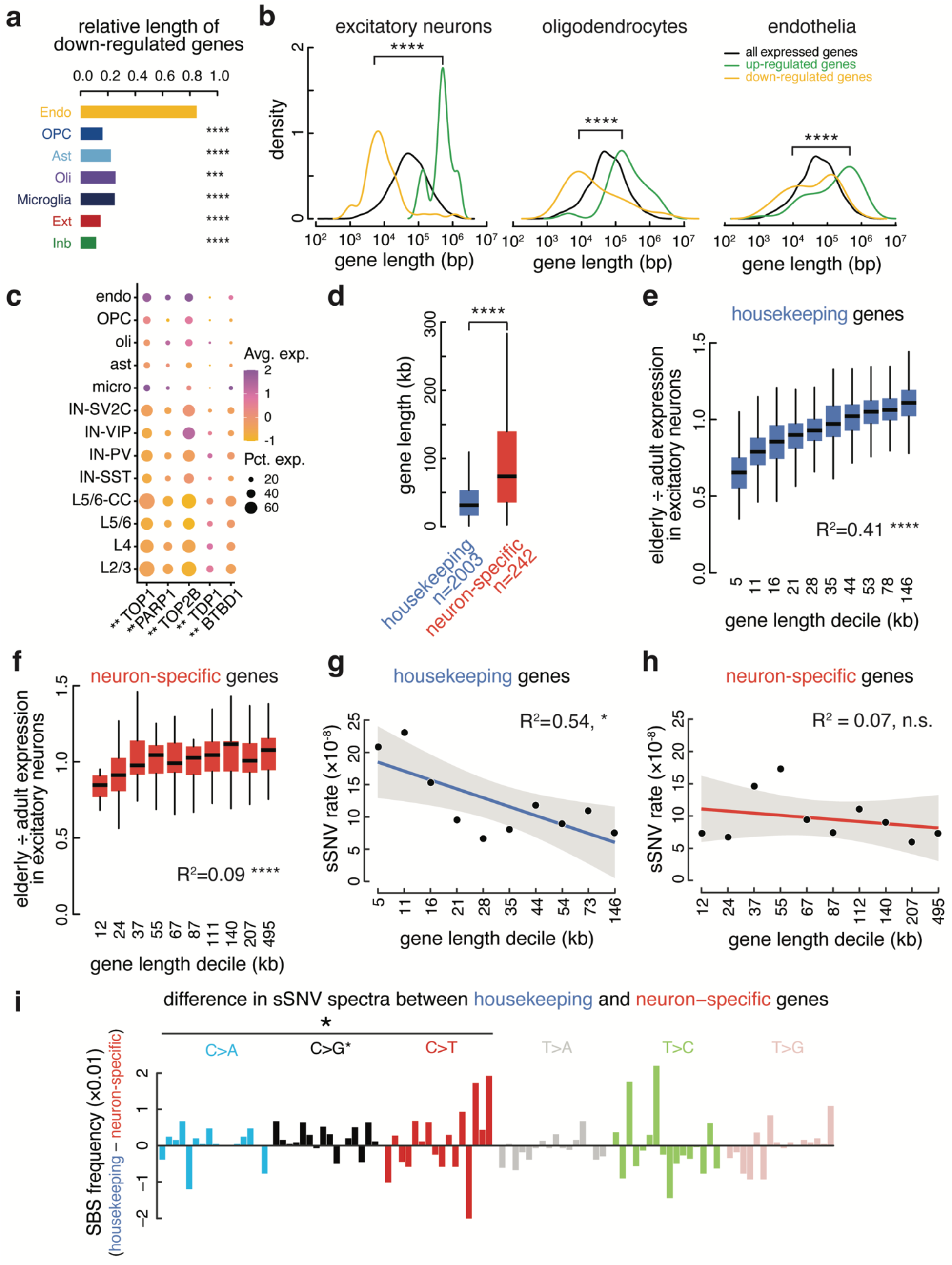
(below) Size-specific down-regulation of housekeeping genes and preservation of cell identity genes. (**a**) Bar graph showing the size of down-regulated genes by cell type relative to the median gene size for each cell type. (Wilcoxon Rank Sum Test). (**b**) Density plots of lengths of all expressed genes (black), upregulated genes (green), and downregulated genes (yellow) in three cell types. (**c**) Expression in topoisomerase complex genes across cell types. Stars denote significant difference between neurons and non-neurons in the percent of cells expressing the gene (Wilcoxon test). (**d**) Housekeeping genes are significantly shorter than neuron-specific genes. (n = number of genes in each category). (**e**) Elderly/adult fold change of housekeeping genes by size decile in excitatory neurons (R^2^ = 0.41). The shortest housekeeping genes have the lowest expression in elderly cells. (**f**) Elderly/adult fold change of neuron-specific genes by size decile in excitatory neurons (R^2^ = 0.09). (**g, h**) Somatic SNV rate per base pair in housekeeping genes **(g)** negatively correlates with gene size, plotted by decile, while **(h)** neuron-specific genes do not show a significant relationship with gene size. 95% confidence interval shown in gray. (**i**) Difference in single-base substitution frequency between housekeeping and neuron-specific sSNV mutation spectra. C>N substitutions are significantly enriched in housekeeping genes, with particular enrichment in C>G substitutions (exact binomial test). Trinucleotide contexts are the same as shown in Extended Data figure 6. (*, p < 0.05; **, p < 0.01; ***, p < 0.001; ****, p < 0.0001). All box plots depict median, and first and third quartile. Whiskers show one and a half times the interquartile range beyond the first and third quartiles.

While in opposition to the relationship observed in many tissues, our data is in agreement with data from mouse cerebral cortex and bulk-sorted retinal ganglion cells, where longer genes increase in expression during aging ^38^. Interestingly, a larger percentage of neurons than non-neurons express the topoisomerases TOP1 and TOP2B, and the topoisomerase interactors PARP1, TDP1, and BTBD1 (p = 1.55 x 10^-3^, Wilcoxon test) (fig. 6c). Neurons rely on topoisomerase activity to mitigate the torsional stress generated when unwinding neuronal genes during transcription^39,40^, which tend to be longer than average^41^ (Figure 6d). As shown in Figure 4 and extended data Figure 5, core metabolic genes were downregulated during aging, while neural cell identity genes were generally preserved. Thus the downregulation of short genes in the aging brain may be a function of the general downregulation during aging of core metabolic genes, which tend to be short, while the relative increase in expression of long genes may reflect the preservation of expression of longer, neural cell identity genes, aided by the expression of topoisomerases.

Our combined single-cell genomic and transcriptomic dataset allowed us to probe the relationship between gene size, genome damage, and age-related expression changes in depth at the single neuron level. Since gene length and gene function are related in the brain, we separately analyzed the relationship between gene length and expression change during excitatory neuron aging in neuron-specific genes and housekeeping genes. Housekeeping genes showed a positive correlation between gene length and expression change in aging (R^2^=0.41, p = 5.94 x 10^-229^) (Figure 6e and Extended Data Fig. 9) such that the shortest genes were the most downregulated while the longest showed no change or slightly increasing in aged cells. This pattern resembled the downregulation of short genes and upregulation of long genes observed in the overall transcriptome (Figure 6b). However, across neuron-specific genes there was a significantly weaker relationship (Fisher’s r-to-z transformation, p-value = 5.55 x 10^-11^) between gene length and expression change in aged brains (R^2^=0.09, p-value = 2.99 x 10^-6^) (Figure 6f and Extended Data Fig. 10). Within gene classes the somatic SNV rate mirrored changes in expression during aging; in housekeeping genes the somatic SNV rate decreased as gene length increased (R^2^ = 0.54, p = 1.54×10^-2^) (Figure 6g), while in neuron-specific genes there was no significant relationship between gene length and SNV rate (R^2^=0.07, p = 0.45) (Figure 6h). These data suggest distinct patterns of DNA damage and repair in housekeeping and neuron-specific genes. Indeed, somatic SNVs in housekeeping genes were enriched for cytosine mutations relative to neuron-specific genes (p = 0.03, exact bionomial test). C>G mutations were significantly enriched in housekeeping genes (p = 9×10^-3^, exact bionomial test) and T**C**C>T**T**C and T**C**T>T**T**T were the most prominently increased cytosine substitutions (Figure 6i). The three stop codons, UAA, UAG, and UGA, are AU-rich (7/9 bases are A or U), so C>T and C>A mutations have the highest potential to generate stopgain mutations. Premature stop codons can result in nonsense-mediated decay of mRNA, which could result in lower transcript abundance in elderly neurons. Thus, gene length, gene function, and genome damage combinatorially impact the transcriptome of the aging brain.

## Discussion

Aging is characterized by a progressive decline in function at the organ, tissue, and single-cell levels. Housekeeping functions were the most commonly enriched gene ontology terms for down-regulated genes, dominating the neurons in particular, while neuron-specific genes remained generally flat during aging with no significant changes in expression. As previously reported, the somatic mutation burden of neurons increased with age. The mutation spectrum matched what we and others have previously seen in neurons. De novo signature analysis revealed two signatures, A1 and A2, characterized by T>C and C>T transitions, respectively, that clustered with known somatic mutation signatures in cancer, SBS5 and SBS30, respectively. The etiology of SBS5 is unknown but has been reported previously by us and others to behave in a clocklike-manner in brain and other tissues ^9,10,23,24^. SBS30 increases in tumors with inactivating mutations in NTHL1, a base excision repair protein ^35^, and we found that NTHL1 expression is significantly reduced during aging in excitatory neurons, pointing to a possible vulnerability in aging that permits Signature A2 mutation accumulation.

The relationship between gene length and the aging transcriptome has long been a point of curiosity in the field but thus far, the relationship has varied across tissues ^36–38^. In our analysis, neuron-specific genes, which tend to be longer than average, show little relationship between expression and size. Conversely, housekeeping genes, across all cell types, show a decrease that is inversely proportional to gene length, i.e. shorter housekeeping genes show the largest decrease in expression in aging. Interestingly, the somatic mutation rate per base pair was highest in the shortest housekeeping genes and mutation rate correlated negatively with size. In neuron-specific genes where there was a weaker relationship between size and expression changes, driven more by increased expression of long genes than decreased expression of short genes, the mutation rate per base pair is not significantly correlated with size. These results identify an explanation for the down-regulation of housekeeping genes in aging and demonstrate the negative impacts of aging-associated somatic mutations. They also suggest the possibility of neuron-specific DNA damage repair pathways that are responsible for maintaining the integrity of key neuron identity genes in long lived cells. As the application of single-cell whole-genome sequencing technologies continue to advance to other cell types in the brain, the relationship between somatic mutations and gene expression during aging can be further elucidated, increasing our understanding of the genomic and transcriptomic landscape in the aging brain.

## Extended Data Figures

**Extended Data Figure 1.**
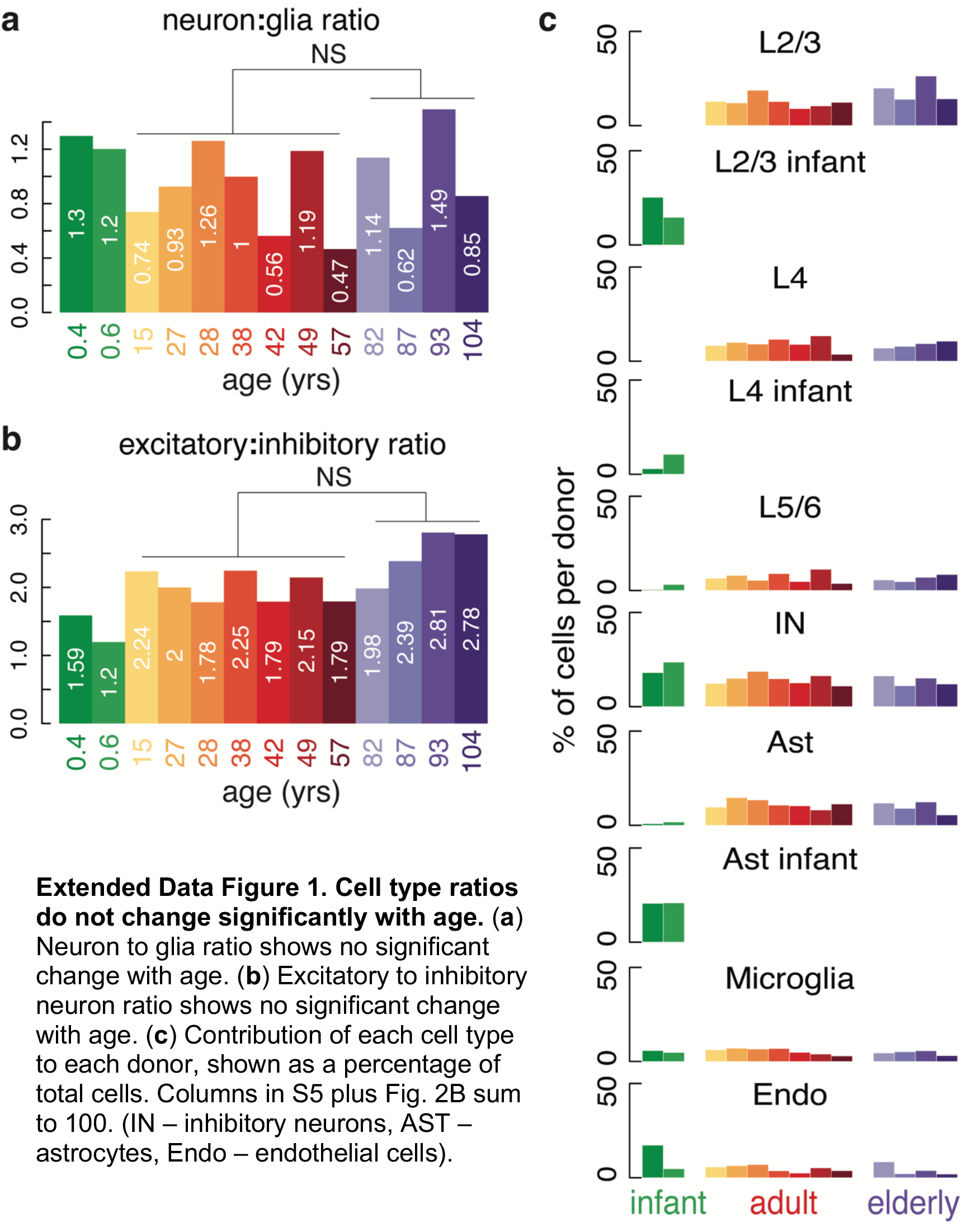
Cell type ratios do not change significantly with age. (**a**) Neuron to glia ratio shows no significant change with age. (**b**) Excitatory to inhibitory neuron ratio shows no significant change with age. (**c**) Contribution of each cell type to each donor, shown as a percentage of total cells. Columns in S5 plus Fig. 2B sum to 100. (IN – inhibitory neurons, AST – astrocytes, Endo – endothelial cells).

**Extended Data Figure 2 (above).**
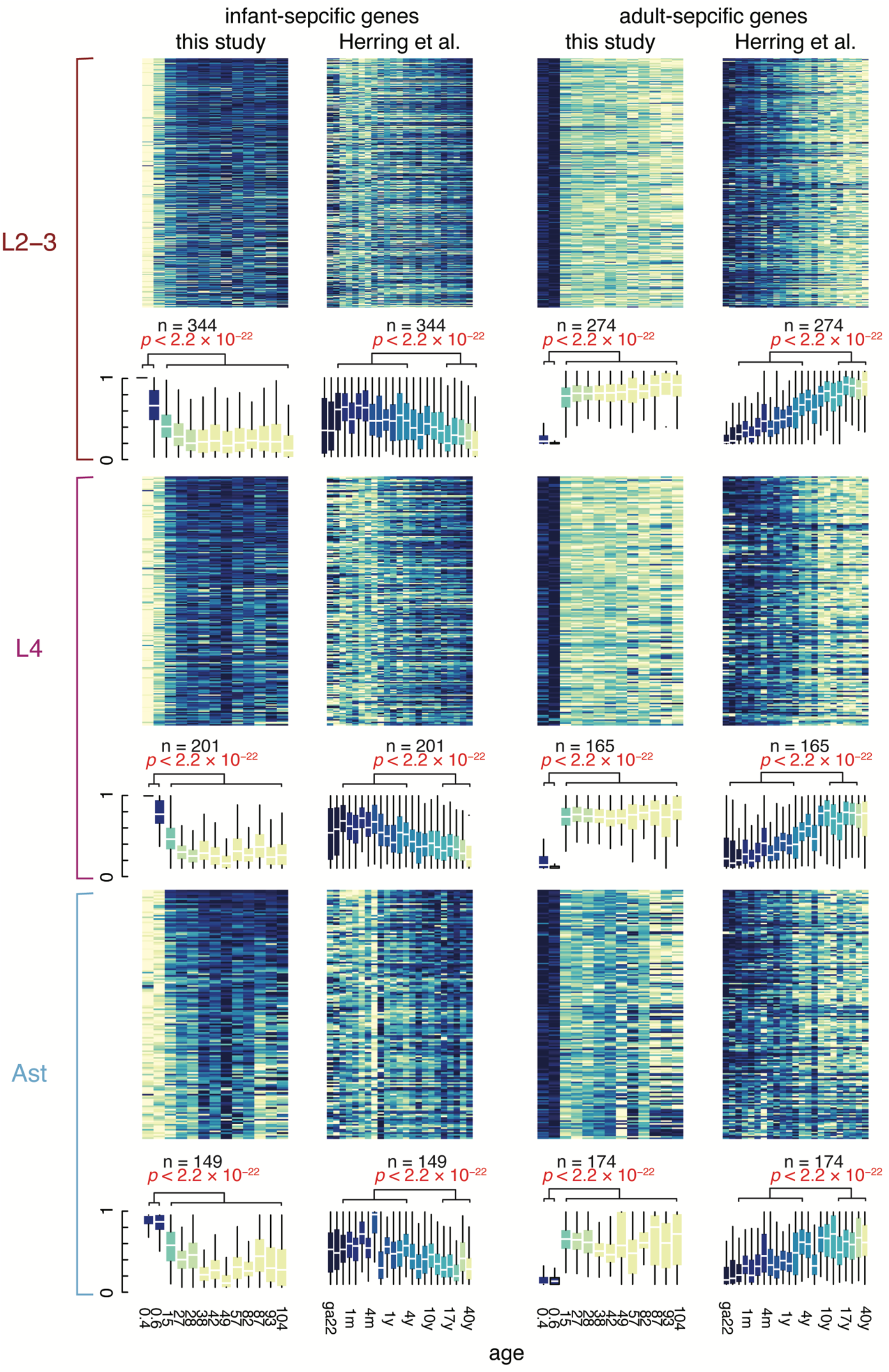
Infant-specific gene expression across ages in two independent data sets. Heatmaps (top) plotting infant-specific gene expression ordered by age in the this study and Herring et al. showing higher expression in infant and gestational cases and lower expression in adults for L2/3 neurons, L4 neurons, and astrocytes. Box plots (bottom) showing mean expression of infant-specific genes and adult-specific genes in L2/3 neurons, L4 neurons, and astrocytes across ages in this study and Herring et al. Expression of infant-specific genes is significantly higher in donors <4 compared to donors >15. Expression of adult-specific genes is significantly lower in donors <4 compared to donors >15. (****, p<0.0001; Wilcoxon Rank Sum test).

**Extended Data Figure 3 (above).**
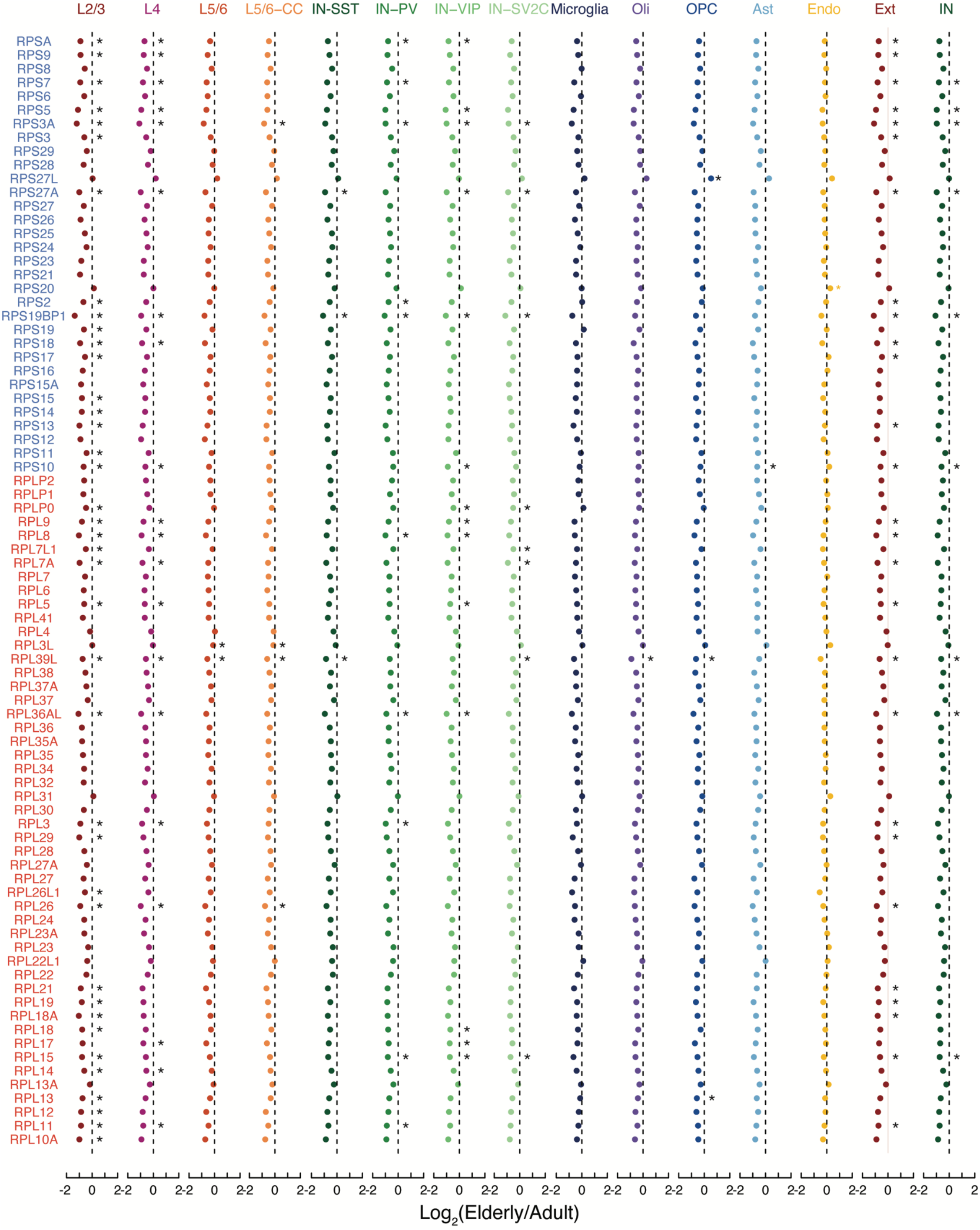
Fold change in elderly vs. adult brains for all ribosomal proteins. Fold change of ribosomal proteins in aging for each cell type. Ext, excitatory neurons; Inb, inhibitory neurons; Oli, oligodendrocytes; OPC, oligodendrocyte precursor cells; Ast, astrocytes; Micro, microglia; Endo, endothelial. (*, p < 0.05; two-sided T-test).

**Extended Data Figure 4.**
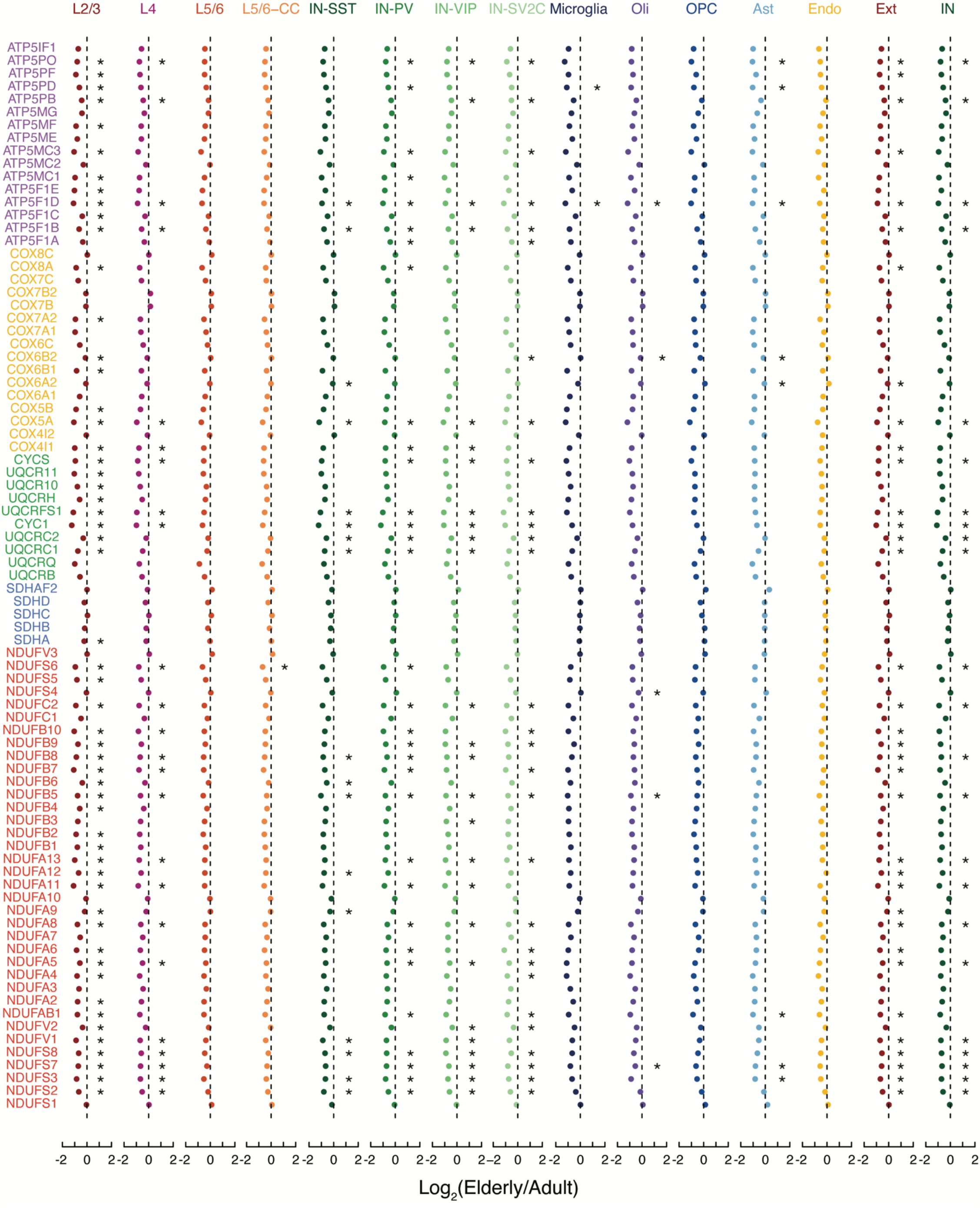
Fold change in elderly vs. adult brains for all nuclear-encoded mitochondrial proteins. Fold change of nuclear-encoded mitochondrial proteins in aging for each cell type. Ext, excitatory neurons; Inb, inhibitory neurons; Oli, oligodendrocytes; OPC, oligodendrocyte precursor cells; Ast, astrocytes; Micro, microglia; Endo, endothelial. (*, p < 0.05; two-sided T-test).

**Extended Data Figure 5.**
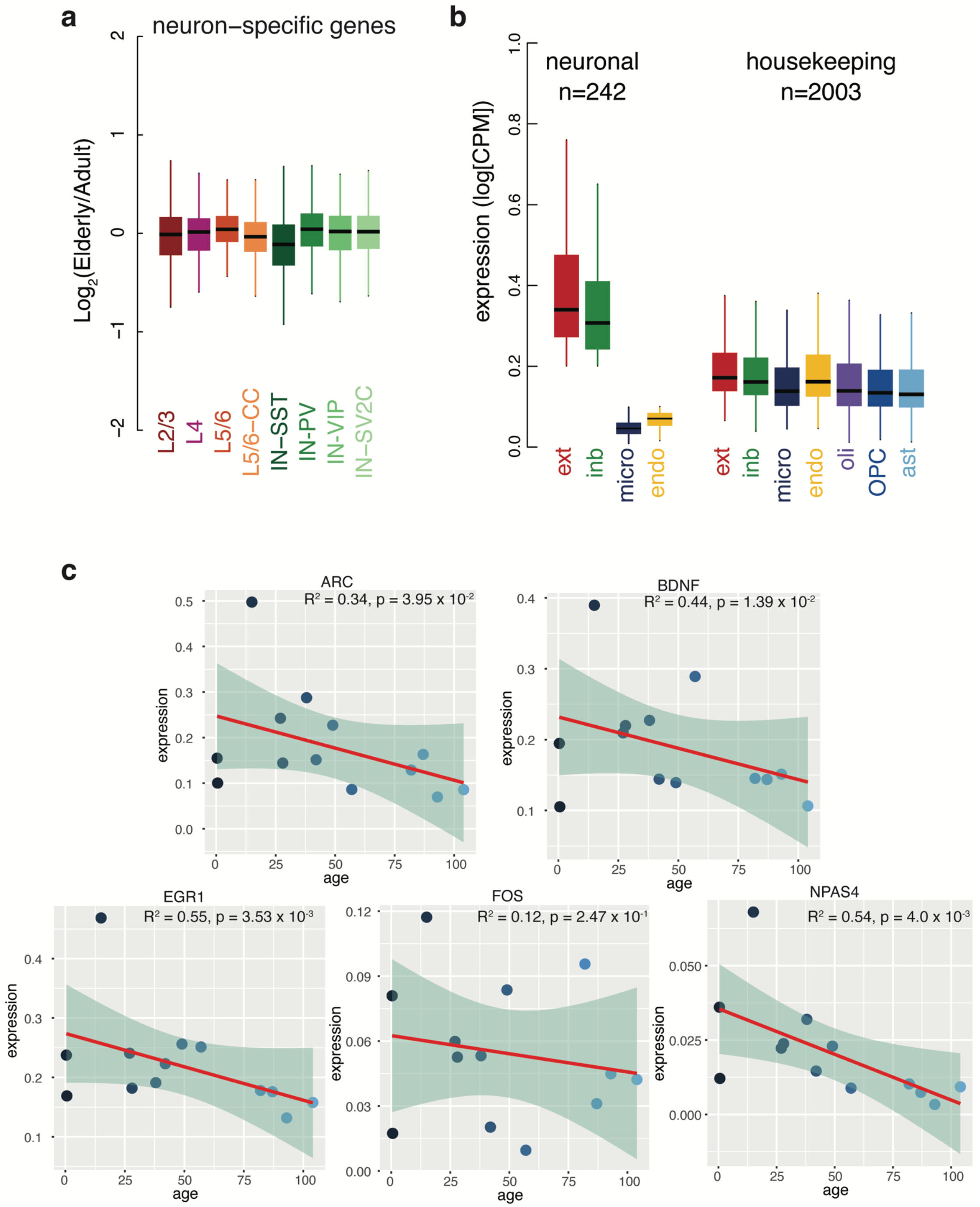
Gene class specific expression and correlation between immediate early gene expression and age (above). (**a**) Expression of neuron-specific genes shows no significant fold change in aging. (**b**) Expression of neuronal and housekeeping genes in the relevant cell types. (**c**) Line represents linear model and shaded area shows a 95% confidence interval. Pearson’s R^2^ and p-value shown.

**Extended Data Figure 6.**
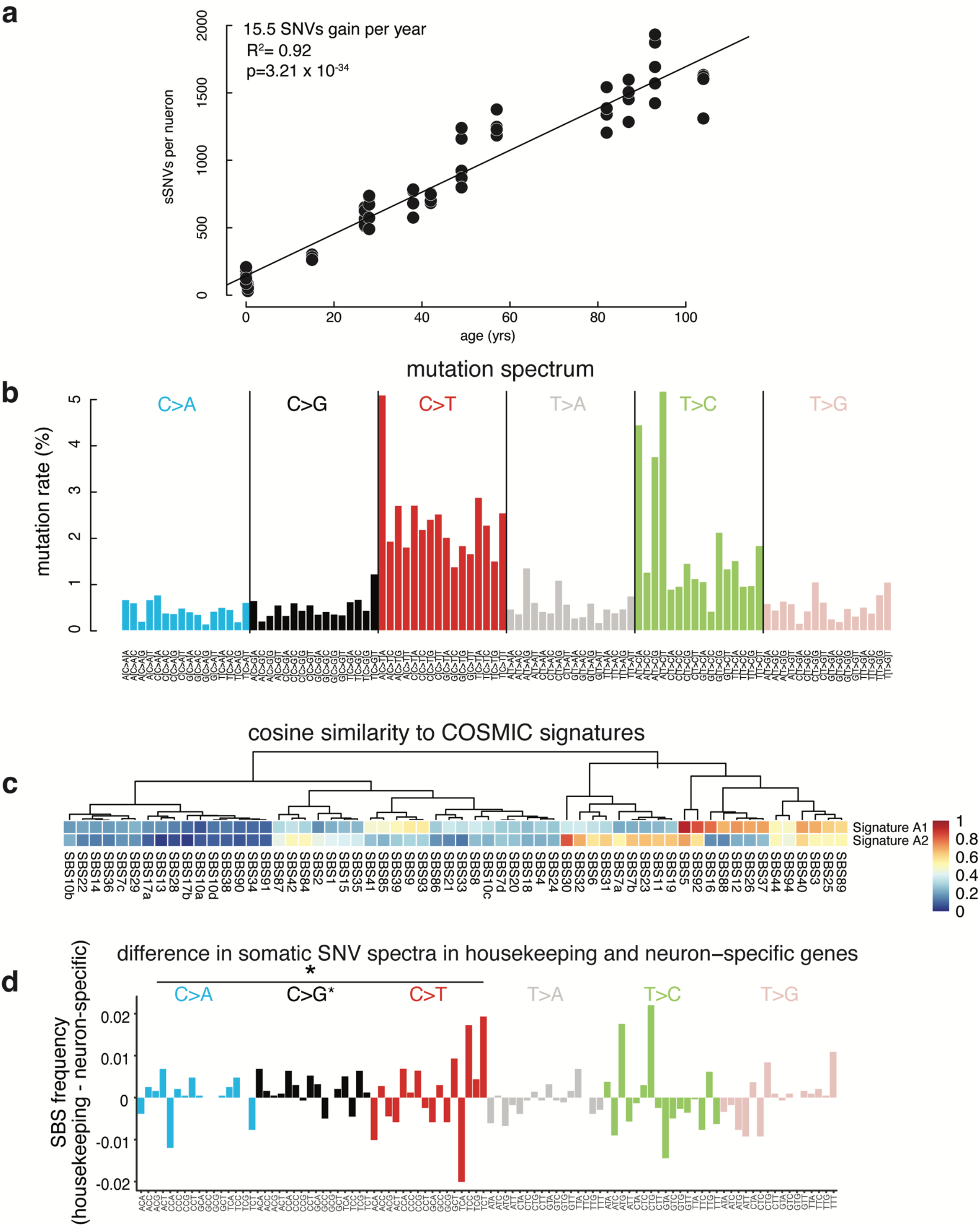
Mutation spectrum of sSNVs in human neurons (above). (**a**) Total mutation accumulation per neuron correlates significantly with age at a rate of 15.5 SNVs gained/year. (**b**) Mutation spectrum of sSNVs called in human neuron scWGS data. Each bar represents a specific mutation in a different trinucleotide context. (**c**) Cosine similarity of the two signatures, A1 and A2, derived de novo from the total mutation spectrum to each single-base substitution signature in the COSMIC data base. Signature A1 is most similar to SBS5. Signature A2 is most similar to SBS30. (**d**) Difference in somatic SNV spectra in housekeeping and neuron specific genes. Same plot as shown in figure 6i but with trinucleotide context specified below each bar.

**Extended Data Figure 7.**
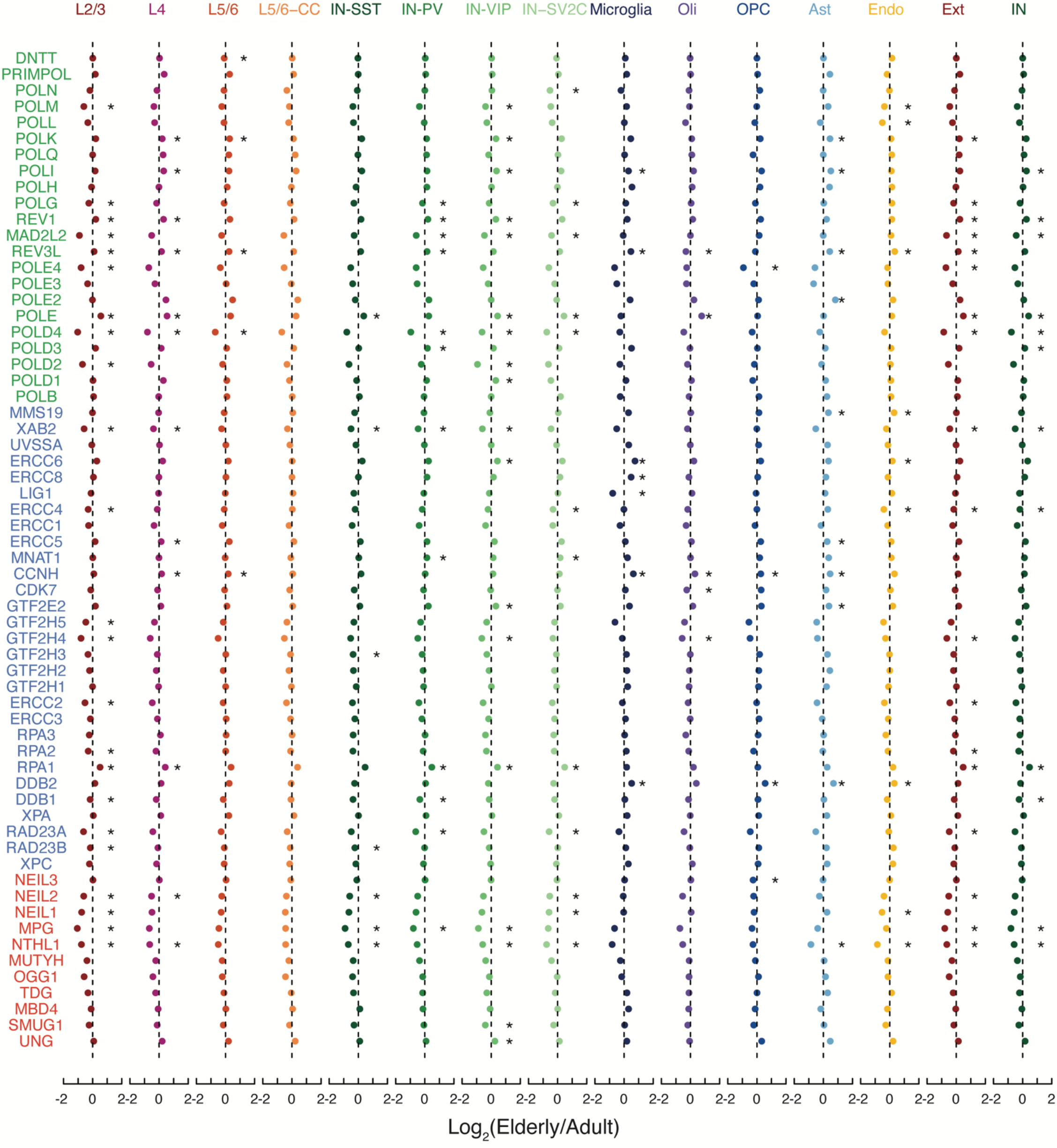
Fold change in elderly vs. adult brains for DNA damage repair genes. Fold change of polymerase genes (green), nucleotide excision repair (blue), and base excision repair (red) in aging for each cell type. Ext, excitatory neurons; Inb, inhibitory neurons; Oli, oligodendrocytes; OPC, oligodendrocyte precursor cells; Ast, astrocytes; Micro, microglia; Endo, endothelial. (*, p < 0.05; two-sided T-test).

**Extended Data Figure 8 (above).**
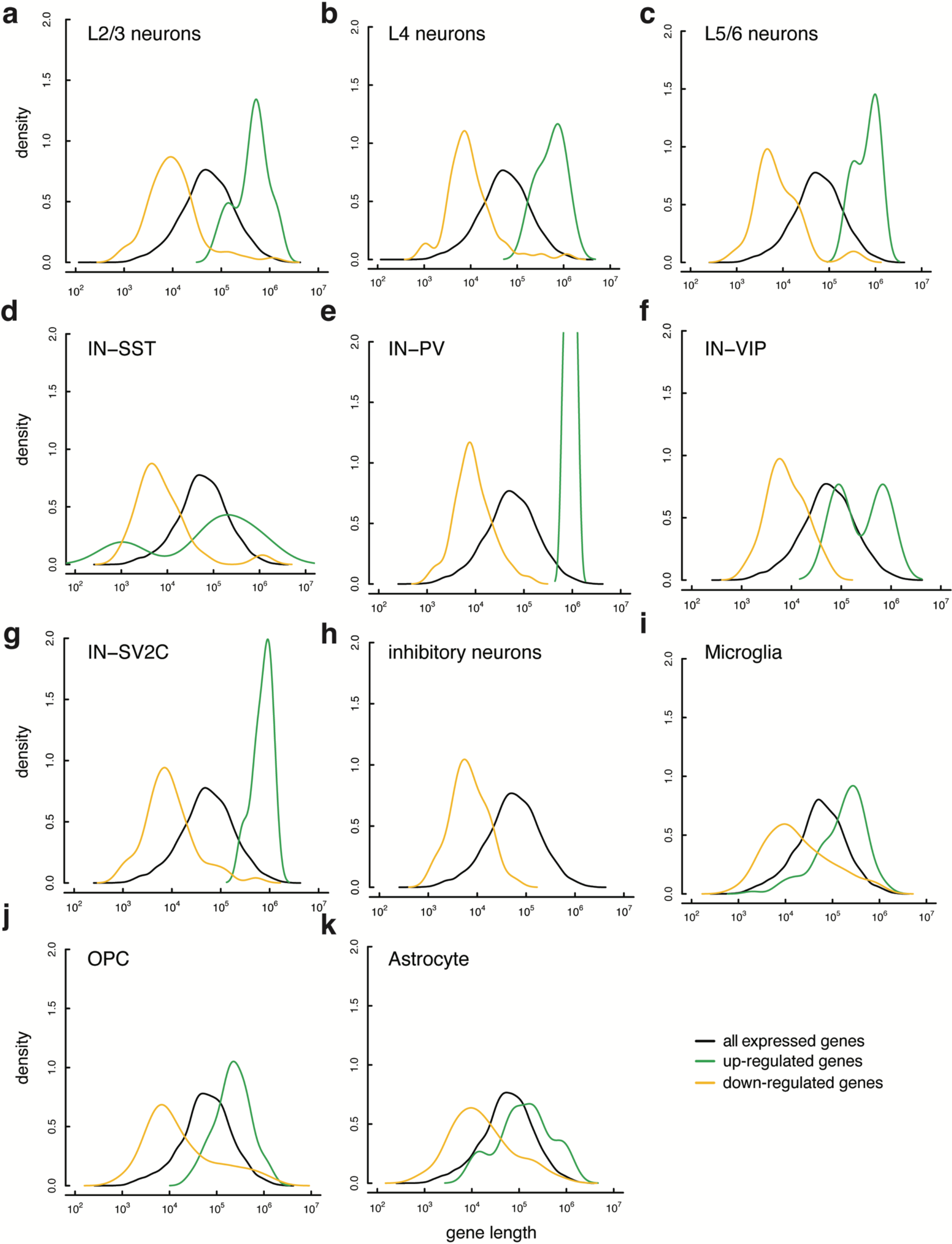
Density plots of gene length for all cell types. Comparison of expressed gene (black), up-regulated gene (green) and down-regulated gene (yellow) size density in (**a**) L2/3 neurons, (**b**) L4 neurons, (**c**) L5/6 neurons, (**d**) inhibitory-SST neurons, (**e**) inhibitory-PV neurons, (**f**) inhibitory-VIP neurons, (**g**) inhibitory-SV2C neurons, (**h**) Inhibitory neurons, (**i**) microglia, (**j**) oligodendrocyte precursor cells, (**k**) astrocytes. L5/6-CC not shown due to low number of differentially expressed genes.

**Extended Data Figure 9 (above).**
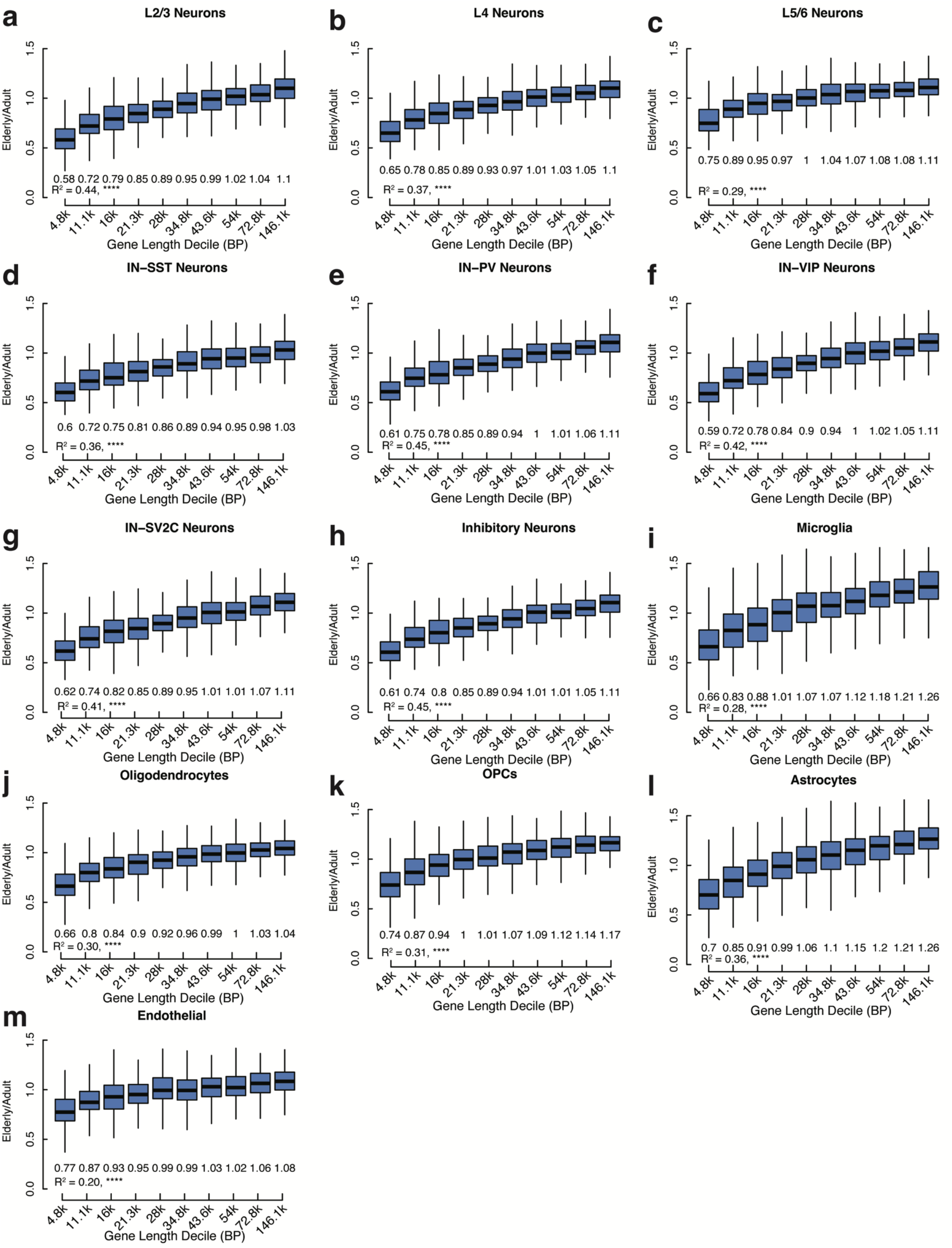
Expression ratio of housekeeping genes by size decile for all cell types. Comparison of elderly to adult expression of housekeeping genes by size decile in (a) L2/3 neurons, (b) L4 neurons, (c) L5/6 neurons, (d) inhibitory-SST neurons, (e) inhibitory-PV neurons, (f) inhibitory-VIP neurons, (g) inhibitory-SV2C neurons, (h) inhibitory neurons, (i) Microglia, (j) oligodendrocytes, (k) oligodendrocyte precursor cells, (l) astrocytes, and (m) endothelial cells. Median fold change shown under each box plot. L5/6-CC neurons not shown due to low number of cells in the cluster. All box plots depict median, and first and third quartile. Whiskers show one and a half times the interquartile range beyond the first and third quartiles.

**Extended Data Figure 10 (above).**
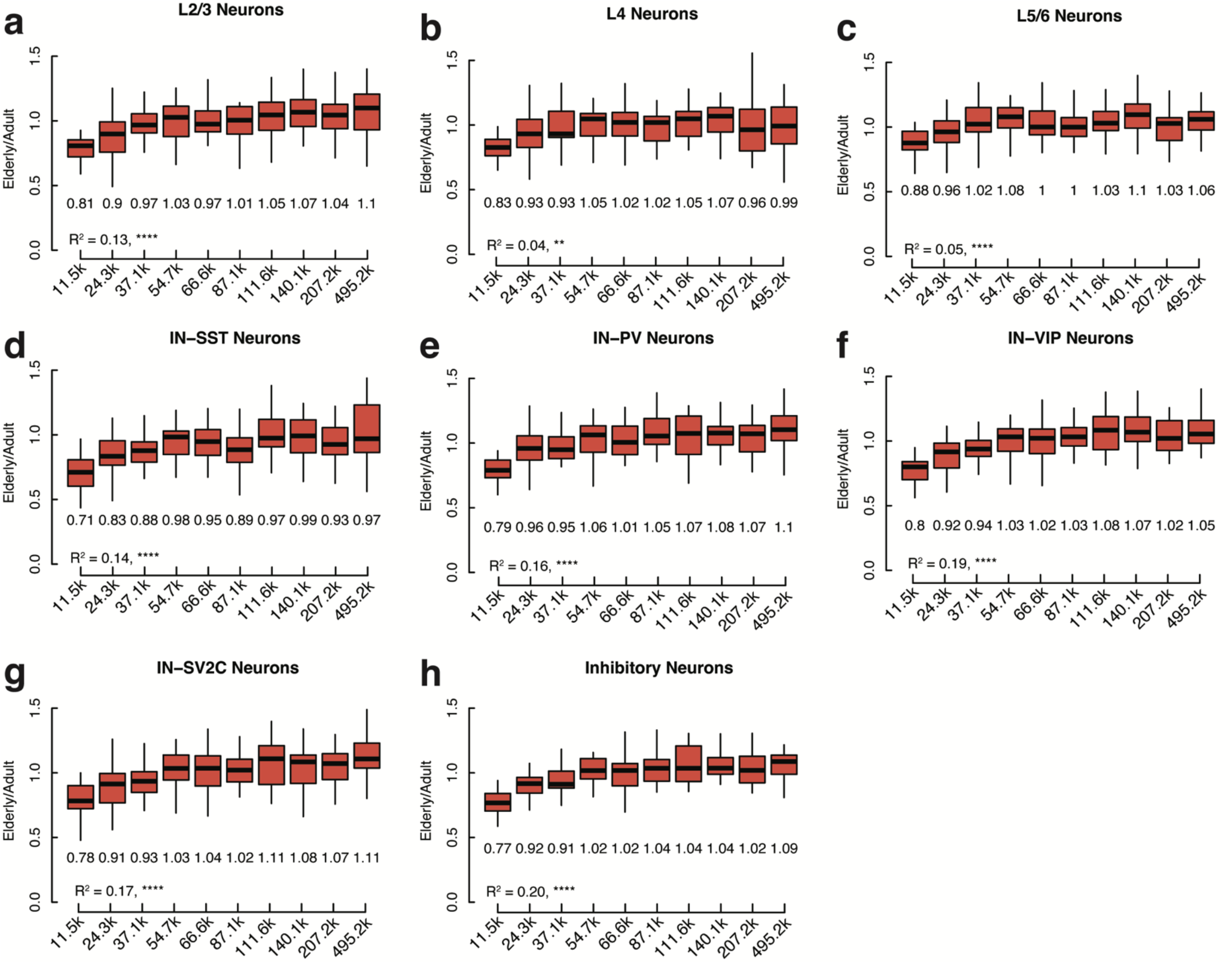
Expression ratio of neuron-specific genes by size decile for all cell types. Comparison of elderly to adult expression of neuron-specific genes by size decile in (a) L2/3 Neurons, (b) L4 neurons, (c) L5/6 neurons, (d) inhibitory-SST neurons, (e) inhibitory-PV neurons, (f) inhibitory-VIP neurons, (g) inhibitory-SV2C neurons, and (h) inhibitory neurons. Median fold change shown under each box plot. L5/6-CC neurons not shown due to low number of cells in the cluster. All box plots depict median, and first and third quartile. Whiskers show one and a half times the interquartile range beyond the first and third quartiles.

## Methods

### Tissue Procurement

All tissue was provided by the National Institutes of Health NeuroBioBank, which obtained written authorization and informed consent for all donors. Cases were selected based on RNA quality, age at time of death, and absence of a history of neurological disease or evidence of neuropathology in the tissue. Brodmann Area 9 or adjacent Brodmann Area 46 of prefrontal cortex was provided for each donor and used for both snRNA-seq and scWGS. To obtain the donor reference genomes, bulk DNA samples were collected from donor-matched tissues, which includes heart, liver, muscle, cerebellum, or cortex. Bulk DNA WGS data for donors 1278, 4638, 4643, 5657, and 5817 (0.4yo M, 15yo F, 42yo F, 82yo M, and 0.6yo M) were obtained from a prior study^1^.

### Nuclei isolation from fresh frozen tissue samples

Nuclei isolation protocol was adapted from two previous publications^2,3^. All procedures were performed on ice or at 4°C. Fresh frozen samples were processed using a 7ml glass dounce homogenizer with approximately 20µg tissue in 5 ml of filter-sterilized tissue lysis buffer (0.32M sucrose, 5mM CaCl_2_, 3mM MgAc_2_, 0.1mM EDTA, 10mM Tris-HCl (pH 8), 0.1% Triton X-100, 1mM fresh DTT). The homogenized solution was loaded on top of a filter-sterilized sucrose cushion (1.8M sucrose, 3mM MgAc_2_, 10mM Tris-HCl (pH 8), 1mM DTT) and spun in an ultracentrifuge in an SW28 rotor (13,300 RPM, 2hrs, 4°C) to separate nuclei.

For nuclei isolated for snRNA-seq, after spinning, supernatant was removed and nuclei were resuspended (1% BSA in PBS plus 25ul 40U/ul RNAse inhibitor), then filtered through a 40µm cell strainer. After filtration, nuclei were counted using trypan blue and an automated hemocytometer (Countess II; Invitrogen) and diluted to a concentration of 1000 cells/µl.

For nuclei isolated for scWGS, supernatant was removed and nuclear pellets were resuspended in ice-cold resuspension buffer (8.5mL 1×PBS/3mM MgCl_2_ + 1mL 1xPBS/3mM MgCl_2_/1% BSA + 500µL Sucrose Cushion), filtered with a 40µm cell strainer, then stained with anti-NeuN antibody (directly conjugated to Alexa Fluor 488; Millipore cat. No. MAB377X, clone A60; 1:1,250) and an anti-rabbit IgG Alexa Fluor 647 antibody as a negative control for 30min. Using a BD biosciences FACSAria Fusion machine and BD FACSDiva Software, forward scatter A (FSC-A) was first used to isolate large non-replicating cells. NeuN staining produced a bimodal signal distribution, distinguishing NeuN^+^ and NeuN^−^ nuclei. Large neuronal nuclei, representing excitatory pyramidal neurons, were further identified by collecting the nuclei with highest NeuN signal among the NeuN^+^ neuronal fraction, and gating for the population with the highest FSC-A signal and excluding Alexa-Fluor-647-high events^4^ (Fig. S7). This non-replicating high-FSC-A plus high-NeuN population was confirmed to comprise an excitatory neuron population, comprising around 10% of the total population of nuclei in each sample^4^.

### Droplet-based snRNA-seq

Droplet-based libraries were generated using the Chromium Single Cell 3’ v3 or v3.1 reagent kits (10x Genomics) according to manufacturer’s instructions. Resulting libraries were indexed with the KAPA Unique Dual-Indexed Adapter Kit (Roche KK8726) and sequenced on an Illumina Novaseq 6000 with 150 paired-end reads by Genuity Sciences (Dublin, Ireland).

### scWGS of neurons using PTA

Single neuronal nuclei, prepared as described above, were whole-genome amplified by primary template-directed Amplification (PTA) ^5,6^ using the ResolveDNA Whole Genome Amplification Kit (BioSkryb Genomics). First, nuclei were sorted into cold 96-well plates pre-loaded with 3µl cold Cell Buffer (BioSkryb) one-per-well. Nuclei were lysed per kit protocol by addition of 3 µl MS mix followed by a brief spin-down, then 1 minute of agitation at room temperature at 1,400 RPM on a plate mixer, then 10 minutes on ice. Next. 3ul SN1 buffer was added to each well and the plate was again spun down and agitated at 1,400 RPM for 1 minute.

Next, 3µl SDX buffer was added, and the plate was again spun and agitated at 1,400 rpm for 1 minute. Then, the plate was incubated at room temperatire for 10 minutes. Next reaction mix + enzyme were added to each well, for a total reaction volume of 20 µl/well. PTA was carried out for 10 hours at 30 °C, followed by enzyme inactivation at 65 °C for 3 min. Amplified DNA was cleaned up using an in-house carboxyl magnetic bead cleanup solution (0.024M PEG-8000, 1M NaCl, 1mM EDTA, 10mM Tris-HCl pH 8, 0.055% Tween 20, and 1.5ml Cytiva Sera-Mag SpeedBeads™ Carboxyl Magnetic Beads, hydrophobic per 50 ml). DNA yield was determined using the QuantiFluor dsDNA System (Promega). Samples were subjected to quality control by multiplex PCR for four genomic loci on different chromsomes. Amplified genomes showing positive amplification for all four multiplex PCR loci were prepared for Illumina sequencing.

Libraries were prepared following a modified KAPA HyperPlus Library Preparation protocol described in the ResolveDNA EA Whole Genome Amplification protocol. In brief, the fragmentation step was skipped and end repair + A-tailing was performed for 500 ng amplified DNA input. Adapter ligation was then performed using the SeqCap Adapter Kit (Roche, 07141548001). Ligated DNA was cleaned up using in-house beads and amplified through an on-bead PCR amplification step. Amplified libraries were selected for a size of 300–600 bp using double-size selection. Libraries were subjected to in-house quality control using a 5300 Fragment Analyzer Bioanalyzer for DNA fragment size distribution (Agilent Technologies). Successfully prepped samples were sent to Genuity Sciences (Dublin, Ireland) for DNA sequencing, who further tested for quality using TapeStation (Agilent Technologies) prior to processing. Single-cell PTA-amplified genome libraries were sequenced on the Illumina NovaSeq 6000 platform (150 bp × 2) at minimum 20× coverage (Table S9)

### Bulk DNA isolation

Genomic DNA was isolated using the Qiagen DNA Mini kit (Qiagen 51304) according to manufacturer’s protocol for tissues. Approximately 25mg of fresh frozen tissue was minced on ice into small still-frozen pieces. Tissue was transferred to a dry-ice chilled sterile 1.5ml microcentrifuge tubes with 180ul of Buffer ATL. Then, 20ul of Proteinase K (20mg/ml) was added prior to 4 hours of agitation at 56 degrees centigrade on a thermomixer (1400 RPM). DNA isolation proceeded as written in the protocol with the inclusion of the optional RNase A treatment step. A small sample was sent for Fragment Analyzing and gDNA quality assessment.

### Bulk DNA library preparation and sequencing

Bulk DNA was isolated as described above and libraries were prepared following the KAPA HyperPlus Library Preparation protocol. The KAPA fragmentation step was included in the bulk processed gDNA samples. Bulk gDNA sample libraries were sent to Genuity Sciences (Dublin, Ireland) and sequenced on the Illumina NovaSeq 6000 platform (150 bp × 2) at minimum 30× coverage and used as a reference genome against the case-match single cell genomes. Bulk DNA for cases 1278, 4638, 4643, 5657, and 5817 was previously isolated and sequenced for Lodato et al. (2018)^1^ on an Illumina Hi-Seq X Ten platform by Macrogen Genomics or the New York Genome Center (New York, NY).

### Analyses of snRNA-seq data

The snRNA-seq reads were aligned to the human genome and assigned to genes (GENCODE v32) via Cell Ranger (v6.0.2) with parameters --expect-cells=10000 --include-introns=true ^7^. The barcode and UMI solved counts were further processed with Seurat ^8^ (4.3.0). The following filtering criteria were applied to each sample and cell: more than 100 cells in the sample, reads from mitochondrially-encode genes less than 5%, and more than 500 expressed gene in the cell. We further filtered samples “5817 200102”, “5288 200128”, and “5887 PFC 210601” according to their poor consistency with other samples in the following clustering results. To minimize false discovery and focus on universal changes in aging, mitochondrially-encoded genes and genes in sex chromosomes were removed in the downstream analysis. The filtered data were log normalized with a factor of 10,000. The top 8,000 variable features were selected for PCA, clustering, and UMAP analysis. The top 30 principal components and 0.5 resolution were used for KNN-graph based clustering, yielding 39 clusters.

Each of the cells in this study was anchored to the cells from Velmeshev et al. using the RPCA method with the top 30 principal components ^8–10^. For each of our 39 clusters, the percentages of cell types according to Velmeshev et al. were calculated, and the dominant cell types were used for each cluster. Those cell types with ambiguous cell types according to Velmeshev et al. were considered as artifacts and removed from the downstream analysis. For analyses where excitatory neuron layer or inhibitory neuron subtype are not specified, layer- and subtype-specific clusters were combined and analyzed as a group. Specifically, all neurons from the L2/3, L4, L5/6, and L5/6-CC clusters were combined into a non-layer-specific group of excitatory neurons and neurons from the IN-SST, IN-SV2C, IN-PV, and IN-VIP clusters were combined into a non-subtype-specific group of inhibitory neurons.

We identified changes in transcript abundance during aging using the default parameters in the Seurat R package for differential expression testing. Genes were expressed in at least 25% of elderly or adult cells and the minimum magnitude of the log_2_ fold change was 0.5. The same process was used to identify genes differentially expressed between infant cells and adult cells.

### Gene Ontology analysis

Gene Ontology analysis of biological processes was performed on the differentially expressed genes for each cell type, both up and down-regulated, using the R package “gprofiler2” with the correction method set to “fdr” and source set to “GO:BP”. For each cell type we used the active genes as the background gene set (indicated on supplementary tables as control genes) and defined active genes as those expressed in more than 25% of the cells. Some cell types yielded no enriched or depleted processes and are not shown. Categorical determination of GO terms was done manually. The corresponding category for each term is listed in the supplemental tables.

### Random permutation test for shared down-regulated genes in elderly cell types

To test if there are significantly more genes down regulated in at least one excitatory neuron, at least one inhibitory neuron, and at least two glial cell types than expected, we performed a random permutation test. We randomly picked the same number of expressed genes to designate as down-regulated for each cell type, using a minimum expression cutoff of 25% of the adult cells and 20% of the elderly cells, and recorded the number of shared genes as the expected value. 1,000 permutations were performed and all of the tests yielded fewer shared genes than observed in our data, generating a p-value of less than 0.001.

### Identification of somatic SNVs in neurons

To identify somatic SNVs, we used both scWGS and corresponding bulk WGS data. scWGS and bulk WGS were first processed accordingly to the GATK best practices^11^. Briefly, reads were aligned to the human genome via bwa mem (v0.7.12) with default parameters. PCR duplicates were then filtered using picard, and the remaining reads were recalibrated via GATK “BaseRecalibrator” and “ApplyBQSR”. Genotypes were then identified via GATK “HaplotypeCaller” and “GenotypeGVCFs”. Finally, somatic SNVs were identified by comparing the scWGS to corresponding WGS from bulk tissues via SCAN2 with the following parameters: -- snv-min-sc-dp 5 --snv-min-bulk-dp 10^6^.

### Signature analysis of somatic SNVs

We performed signature analysis for somatic SNVs using the R package MutationalPatterns (v3.10.0)^12^. We first calculated the spectrum of sSNVs in the 96-trinucleotide contexts for each neuron from all donors. A non-negative matrix factorization (NMF) was applied to the spectrum of sSNVs and the signatures were identified. After applying different numbers of signatures in the practice, ranging from one to eight, and found two signatures yield the best performance regarding stability and reconstruction errors (Fig. S5). The signatures (A1 and A2) were then compared to the COSMIC v3 signatures, and cosine similarities between signatures were calculated. To confirm the reproducibility of our signature analysis, a second method SignatureAnalyzer was used with default parameters. SignatureAnalyzer identified similar signatures as MutationalPatterns (Fig. S6).

### Enrichment of somatic SNVs at genes and intergenic regions

To calculate the enrichment of somatic SNVs in genes and intergenic regions, we first simulated random controls with the same mutation spectrum as somatic SNVs restricted to suitable regions (i.e. with enough depth) in our scWGS and bulk WGS dataset. The number of somatic sSNVs and random controls at genes and intergenic regions were then calculated. NMF was further applied to somatic SNVs and random controls at genes and intergenic regions to trace the contribution of signature A1 and A2. Genes were divided into five groups according to their transcriptional activity (CPM) in neurons and glia cells from our snRNA-seq data. The strand for each type of genic somatic SNVs (T>A, T>C, T>G, C>A, C>T, C>G) were further recorded.

### DEPMAP analyses of cell viability impact of upregulated and downregulated genes

The requirement of each gene in overall cell viability was determined using the ‘CancerDependency Map (DEPMAP; version Public 22Q4)’, which provides the cell viability effect of each gene KO across 1078 cancer cell lines of varying origin (PMID: 29083409). Specifically, cell viability is determined via conducting whole-genome pooled CRISPR screening across each cell line (1078 total), based on fold change in abundance of cells harboring Cas9 and guides against each specific gene. For example, if cells transduced with Cas9 and guides against a particular gene have been depleted following the screen, this would indicate an essential gene. The overall impact of gene KO for a given cell line is quantified via a cell population dynamics model called ‘Chronos’ ^13^, which incorporates efficacy of each guide and copy number correction (CRISPR toxicity unrelated to gene function can occur when high copy numbers are subjected to CRISPR mediated strand breaks), to provide an overall ‘gene effect score’ which indicates the probability that a given cell line is dependent on the gene for survival ^14^. Importantly, a value of -1.0 corresponds to the median gene effect score of all common essential genes, while a cell line is considered dependent if the gene effect score is <=-0.5. Positive values would indicate increased cell viability or proliferation following loss of the gene.

Among the upregulated and downregulated ‘hits’ from the scRNA-SEQ, those encoding lncRNAs, ncRNAs, or pseudogenes are not covered in the DEPMAP essentiality analyses and thus were not analyzed for effects on gene viability. Likewise, several coding genes (CECR, NEFL, FTH1, COX4I1, SH3RF3, BMP2K, SHISA8, MYRFL, RPS3A) did not have CRISPR screen data yet available, and were not analyzed.

### Definition of housekeeping and neuron-specific genes

We first calculated the average logged CPMs for each gene in excitatory neurons, inhibitory neurons, microglia, and endothelia. Then we defined housekeeping genes as genes with a difference of less than 0.1 between the four cell types that also had an average logged CPM greater than 0.1 in each cell type. The genes that fit this criteria also have an average logged CPM greater than 0.1 in oligodendrocytes, OPCs, and astrocytes as well. The neuron-specific genes were defined as those genes with average logged CPMs higher than 0.2 in both neuron groups and lower than 0.1 in microglia and endothelia.

## Data availability

Raw RNA-seq and scWGS sequencing data are available at the dbGaP accession (forthcoming). We are currently in contact with dbGaP and all data will be available before publication. An interactive genome browser of the snRNA-seq expression data can be found at: https://genome.ucsc.edu/s/yutianxiong/Weng_Lodato_Aging. The previously published Velmeshev *et al*. data used for cell type annotation was downloaded from SRA accession number PRJNA434002.

## Acknowledgements

We thank the UMass Chan Medical School Flow Cytometry Core and their funding NIH S10 OD028576, as well as T. Krumpoch and S. Pechhold for assistance with single nuclei sorting. We thank D. Kim for manuscript feedback, and T. Chittenden, J. Lopez, R. Li, and A. McLean for their assistance with high throughput sequencing and helpful discussion. Human tissue was obtained from the NIH NeuroBioBank at the University of Maryland and The Sepulveda Research Corporation, and we thank the donors and their families for their generous contributions to research. M.A.L. was supported by R00AG054748, R56 AG078453, The American Federation for Aging Research and The Glenn Foundation for Medical Research, and the Charles H. Hood Foundation. Z.W. was supported by U24HG012343.

## Author Contributions

A.M.J. and M.A.L conceived and designed the study. A.M.J performed single-nucleus RNA-sequencing. T.Y. performed bioinformatic analysis. J.S.Z and A.K.T performed single-neuron sorting and sequencing. Y.K. and A.M.J performed data analysis. A.M.J, and M.A.L wrote the manuscript. M.A.L and Z.W supervised the study.

## Supplementary Figures

**Fig. S1.**
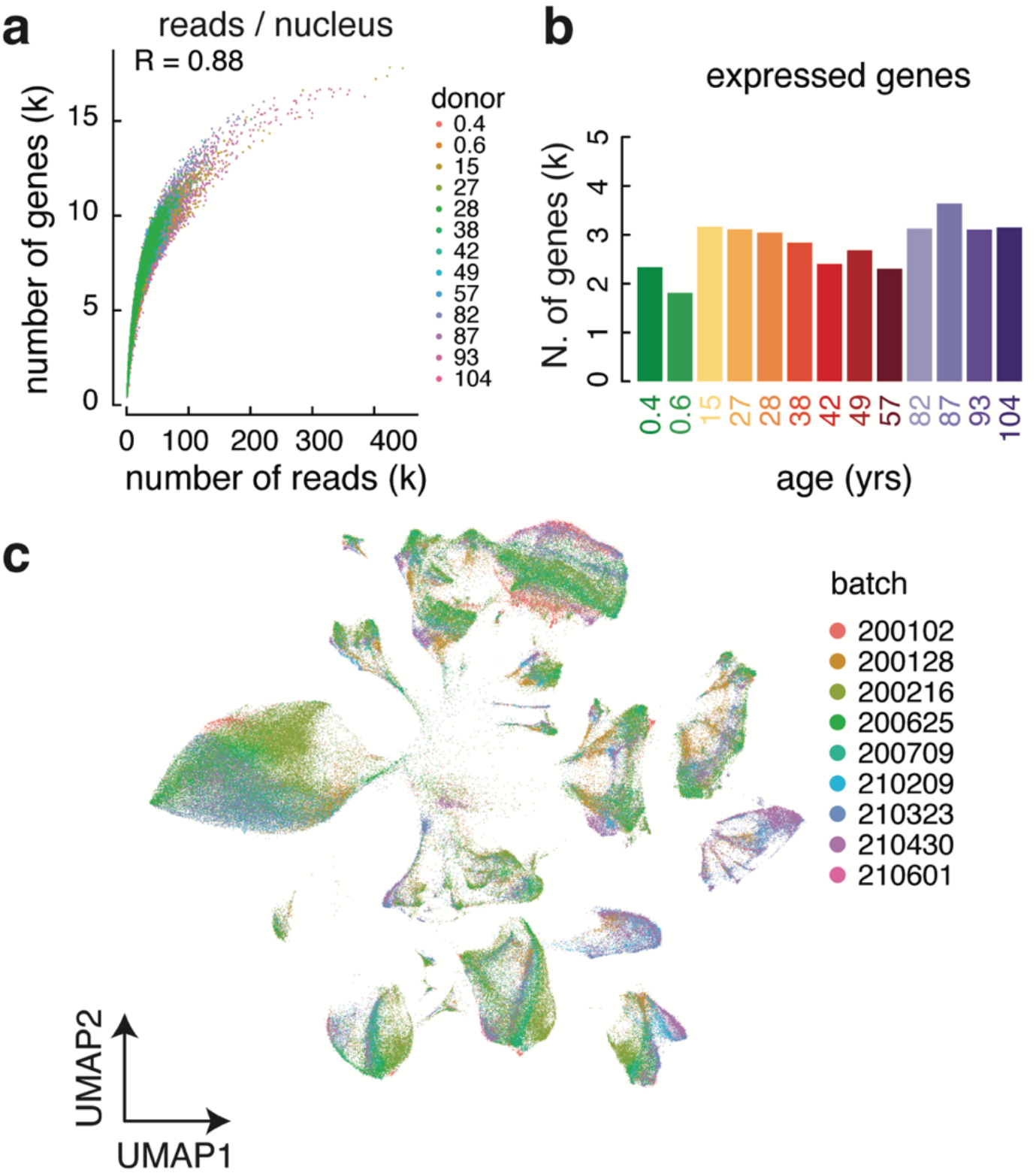
Single-nucleus RNA-seq of the aging human brain. (A) Scatter plot showing the correlation between the number of expressed genes, in thousands, and the number of reads, in thousands, per nucleus, colored by donor. (B) The number of unique genes expressed in each donor did not vary significantly by age. (C) UMAP colored by preparation batch shows uniform distribution of batches across clusters.

**Fig. S2.**
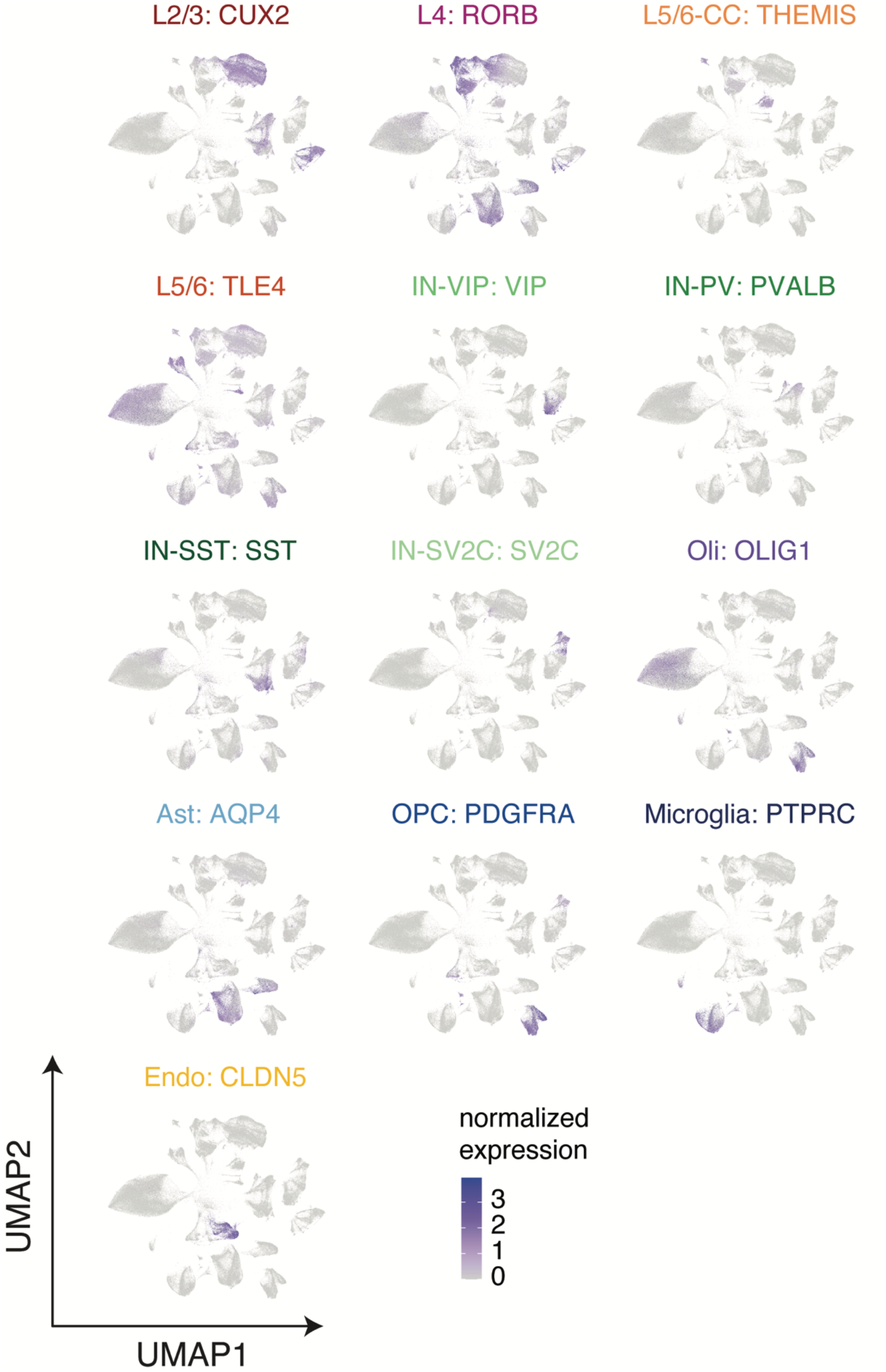
Expression of Marker Genes. Expression of the canonical marker for each cell type is isolated to the corresponding cluster(s) on the UMAP demonstrating cell-type specificity of the clustering.

**Fig. S3.**
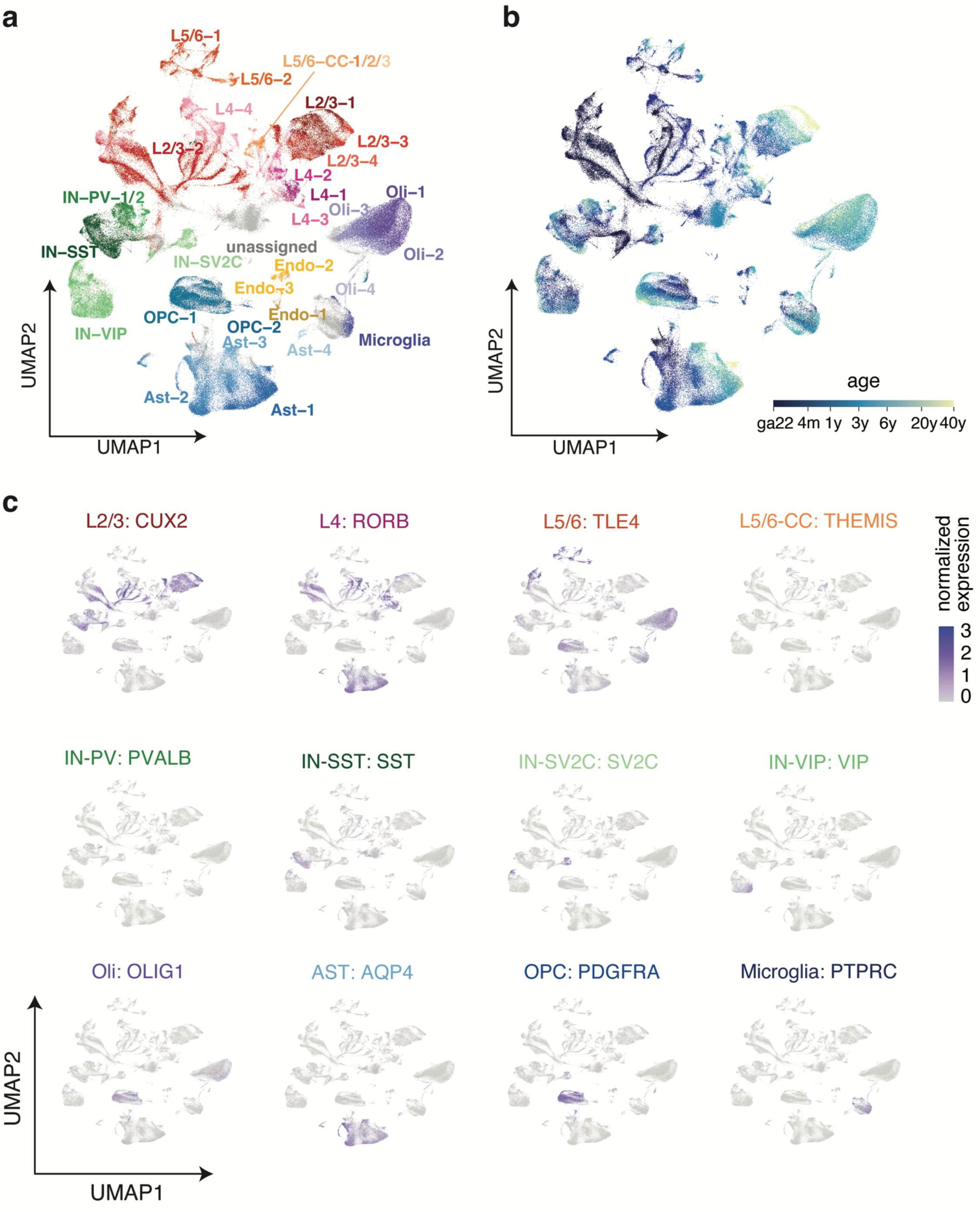
Clustering of Herring et al. Data (above). (A) Donor contribution by age to each cell type. (B) Clustering of Herring et al. publicly available raw data yielded the same sub-clusters as our own data. (C) The same Herring et al. UMAP colored by age of donor reveals the same infant-specific sub-clusters for L2/3 neurons, L4 neurons, and astrocytes. Expression of the canonical marker for each cell type on the Herring et al. data is isolated to the corresponding cluster(s) on the UMAP demonstrating cell-type specificity of the clustering.

**Fig. S4.**
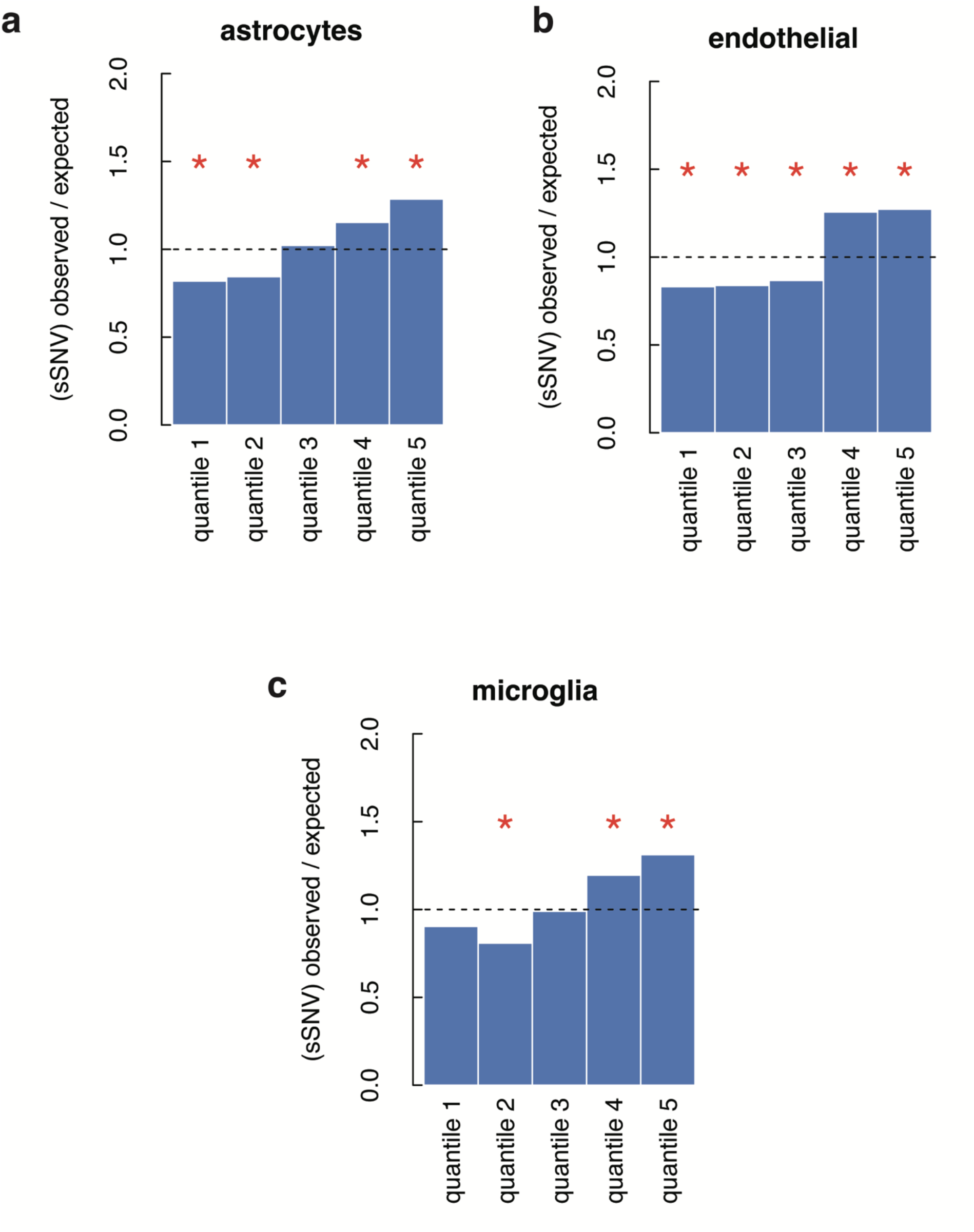
Signature A1 mutation enrichment by expression level. Enrichment of Signature A1 mutations in (A) astrocytes, (B) endothelial cells, and (C) microglia correlates with gene expression. There are more Signature A1 mutations in the highest expressed genes within these cell types than expected. (x^2^-test; *, p < 0.05).

**Fig. S5.**
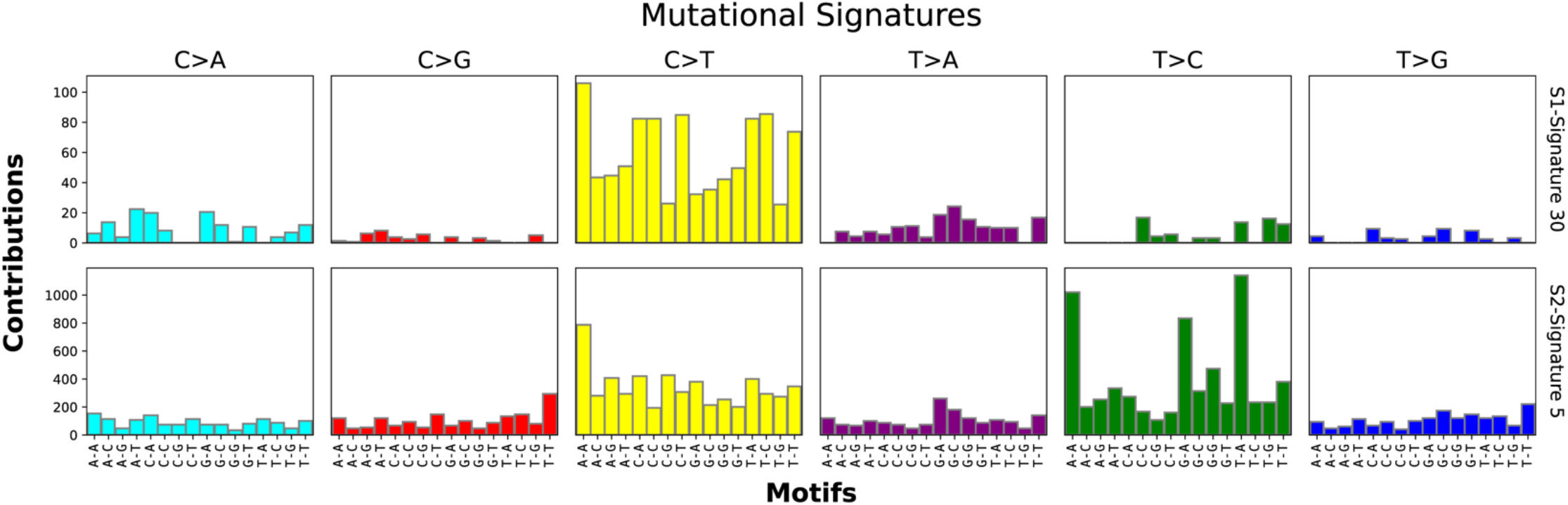
De novo mutation signature analysis by python package signatureAnalyzer. Consistent with the results from MutationalPatterns, two signatures were predicted. The first signature is dominated by C>T mutations, consistent with signature A2, and the second is dominated by T>C mutations, consistent with signature A1.

**Fig. S6.**
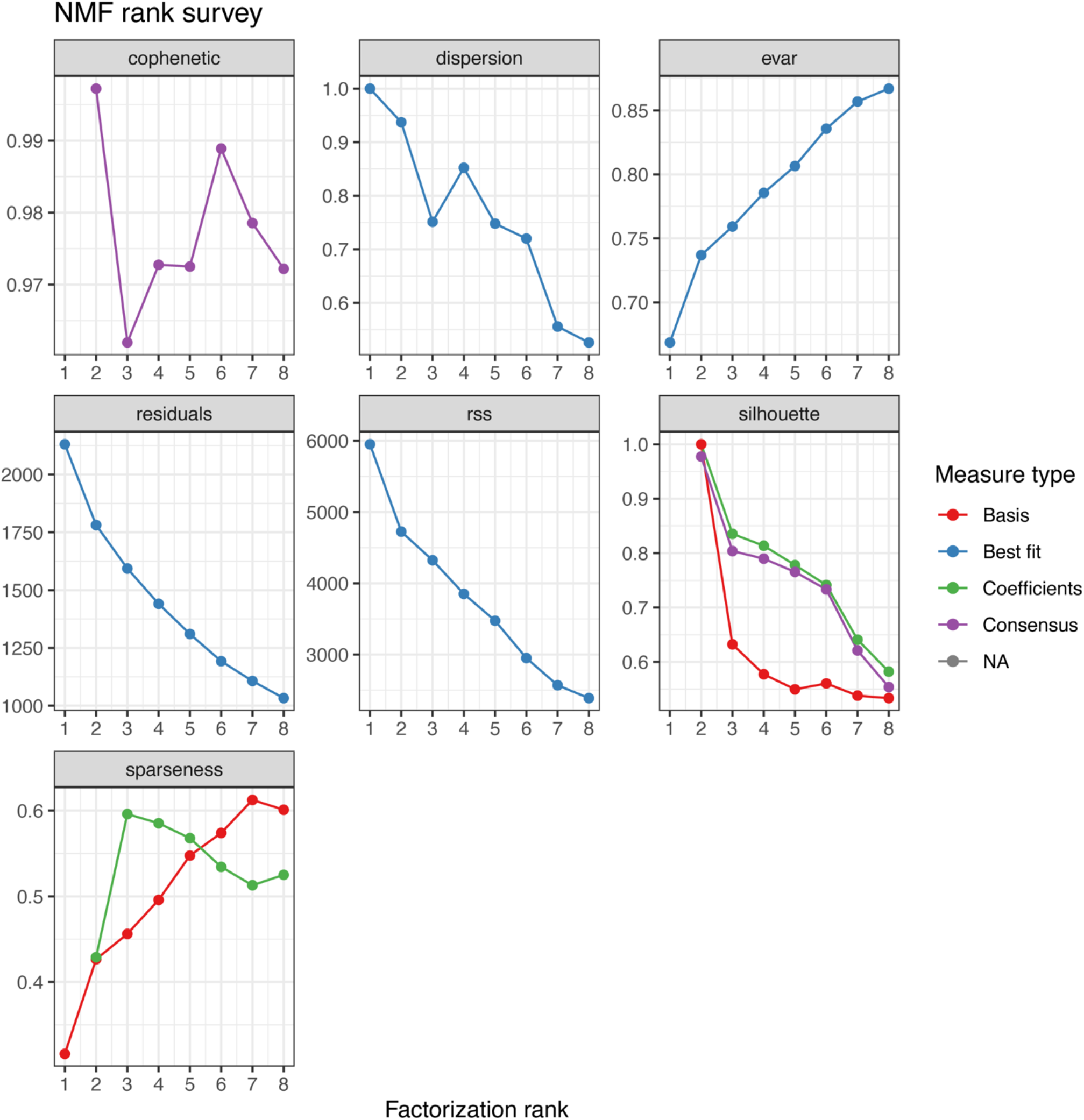
Signature metrics for de novo mutation signature analysis by MutationalPatterns. Mutational signatures of sSNVs are fitted with a NMF-based framework. According to the metrics, we concluded two is the number of signatures that maximize the cophenetic of the decomposition.

**Fig. S7.**
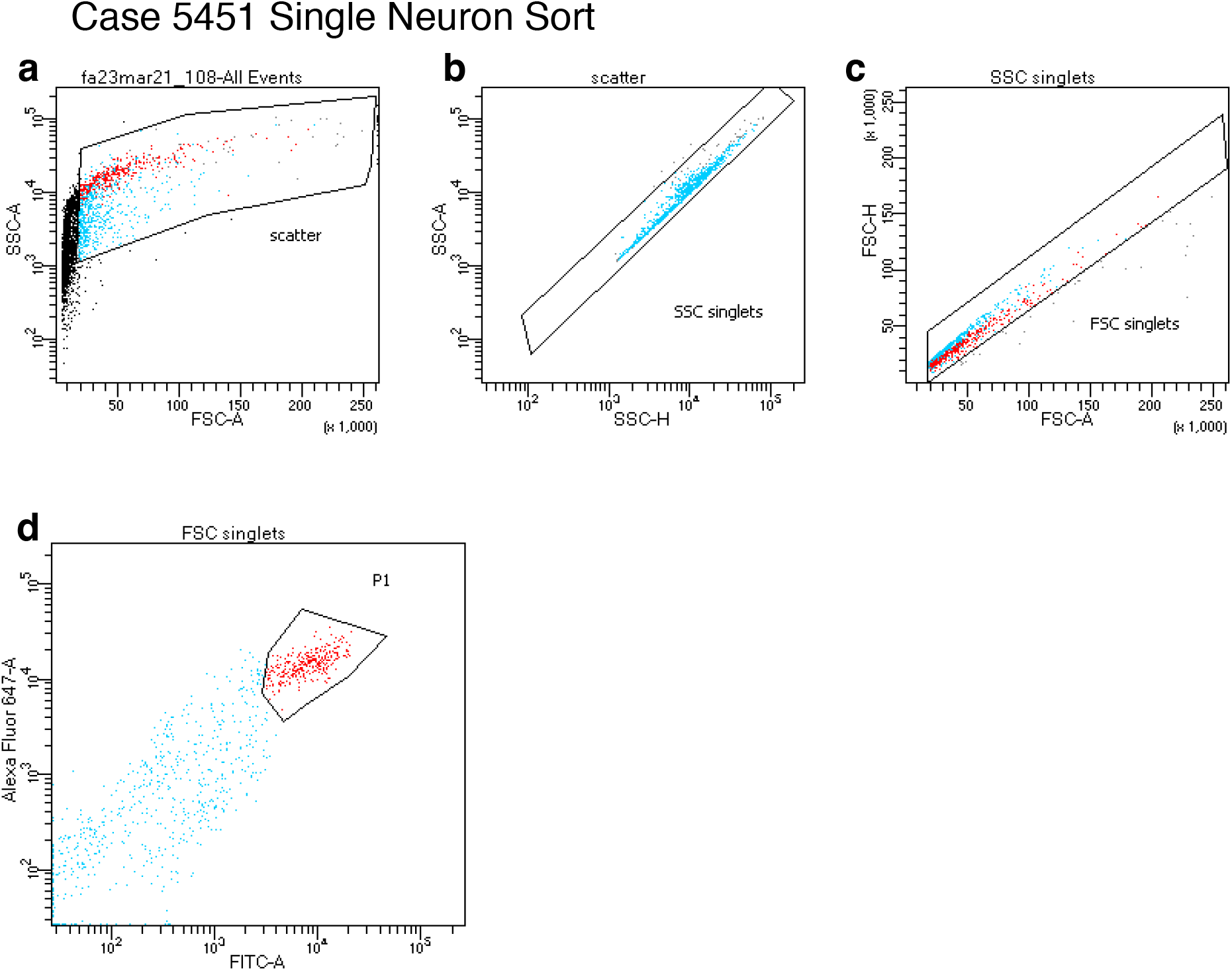
Representative FACS plot for single neuron sorting. (**a**) Gating used to isolate nuclei from debris. (**b, c**) Gating used to remove doublets. (**d**) Gating used to identify neuronal nuclei.

## Supplementary Table Legends

**Table S1. Extended Donor Information.** Case ID, sex, age at death, number of snRNA-seq nuclei, number of scWGS nuclei, post mortem interval (PMI) in hours, cause of death, RNA integrity number (RIN), years in storage, and brain bank information for each donor.

**Table S2. Table of marker genes distinguishing each cluster.** Gene name, log_2_ fold change of expression in marker cluster vs. other clusters, percent of cells in the marker cluster expressing the gene, percentage of cells in other clusters expressing the gene, p-value, q-value, and cluster identity.

**Table S3. Table of cell counts in each cluster broken out by donor.** Each column is a cluster and each row is a donor.

**Table S4. Infant-specific differentially expressed genes.** Table listing genes differentially expressed between infant-specific clusters and adult clusters of the same cell type. Negative log_2_ fold change indicates more expression in adult clusters. Positive log_2_ fold change indicates more expression in infant clusters.

**Table S5. Enriched GO terms for infant specific clusters.** Term name, false discovery rate (FDR), fold enrichment, number of control genes, number of differentially expressed genes number of differentially expressed genes in the term, category represented in figure 2C, and gene hits for each GO germ. Up-regulated indicates the genes that have higher expression in the infant-specific cluster and down-regulated indicates genes that have higher expression in the adult sub-clusters of that type.

**Table S6. Differentially expressed genes common to multiple classes of cells.** Genes differentially expressed across cell types. All genes were down-regulated in elderly brains in all cell types. Excitatory and inhibitory neuron hits indicate the number of excitatory or inhibitory neuron cell types for which the gene was differentially expressed. Glial hits includes endothelial cells along with all glia.

**Table S7. Elderly vs. adult differentially expressed genes.** Table listing genes differentially expressed between elderly cases and adult cases. Negative log_2_ fold change indicates more expression in adult clusters. Positive log_2_ fold change indicates more expression in adult clusters.

**Table S8. Enriched GO terms for elderly vs. adult differentially expressed genes.** Term name, false discovery rate (FDR), fold enrichment, number of control genes, number of differentially expressed genes number of differentially expressed genes in the term, category represented in figure 4A, and gene hits for each GO germ. Up-regulated indicates the genes that have higher expression in elderly brains and down-regulated indicates genes that have higher expression in the adult brains.

**Table S9. Sequencing statistics.** Tabs show sequencing statistics for single cells and associated bulk tissues.

**Table S10. Somatic single nucleotide variation calls.** Summary tab provides SCAN2 sensitivity information for each single cell. Detailed information provides the annovar annotation for each SNV identified with SCAN2.

